# A primate model organism for cardiac arrhythmias identifies a magnesium transporter in pacemaker function

**DOI:** 10.1101/2025.05.28.655959

**Authors:** Stephen Chang, Caitlin J. Karanewsky, Jozeph L. Pendleton, Lu Ren, Aude Anzeraey, Victor Froelicher, David Liang, Andriamahery Razafindrakoto, Hajanirina Noëline Ravelonjanahary, Megan A. Albertelli, Thomas Quertermous, Patricia C. Wright, Martine Perret, Jérémy Terrien, Fabienne Aujard, Joseph C. Wu, Mark A. Krasnow

## Abstract

Cardiac arrhythmias afflict tens of millions of people, causing one-fifth of all deaths^1^. Although mouse models have aided understanding of some pacemaker genes and arrhythmias^2,3^, mice are not known to naturally acquire arrhythmias, and the substantial differences between mouse and human cardiac anatomy and physiology have limited their utility in preclinical studies and pharmacological testing^2,4^. To establish a primate genetic model organism for arrhythmias, we carried out an electrocardiographic (ECG) screen of over 350 lab and wild mouse lemurs (*Microcebus spp.*), an emerging model organism that is among the smallest, fastest-reproducing, and most abundant primates^5^. Twenty-two lemurs (6.2%) were identified with eight different naturally-occurring arrhythmias resembling human ECG pathologies (SSS, PACs, Afib, PVCs, NSVT, STD, iTWs, STE). Pedigree construction showed two were familial, premature atrial contractions (PACs)/atrial fibrillation (Afib) and sick sinus syndrome (SSS), an episodic bradycardia. Genome sequencing of the SSS pedigree mapped the disease locus to a 1.4 Mb interval on chromosome 7 and supported autosomal recessive Mendelian inheritance. The most appealing candidate gene in the interval was *SLC41A2*, a little studied magnesium transporter^6,7^. SLC41A2 is expressed in human iPSC-derived sinoatrial node cells (iSANC) and localizes to the sarcoplasmic reticulum. Although mouse *SLC41A2* knockouts do not show a cardiac pacemaker phenotype^8^, CRISPR-mediated *SLC41A2* knockout altered human iSANC magnesium dynamics and slowed their calcium transient firing rate. The results suggest *SLC41A2* functions cell autonomously and primate-specifically in cardiac pacemaker cells, and that intracellular magnesium dynamics have a crucial but previously unappreciated role in setting pacemaker rate. Thus, mouse lemur is a valuable model for discovering new genes, molecules, and mechanisms of the primate pacemaker, and for identifying novel candidate genes and therapeutic targets for human arrhythmias. The approach can be used to elucidate other primate diseases and traits.

## INTRODUCTION

Systematic genetic studies of a small number of phylogenetically diverse organisms have transformed our understanding of biology and medicine^9–11^. Over the past several decades, mouse has emerged as the biomedical supermodel because of its short generation time for a mammal, ease in lab husbandry, and the ability to create gene knock-out^12,13^, knock-in^14^ and transgenic^15^ mice that model human disease mutations^16^. However, with the expanding use of mouse models has come a greater appreciation and concern that the differences in anatomy, physiology, genetics, and biochemistry between mice and humans limit their utility for the study of many human genes and diseases, and for preclinical testing of therapeutics^5,17–20^. Because of their greater similarity to humans, non-human primates are used in many areas of medical research and preclinical testing. However, ethical and practical considerations such as their large size, great cost, and long maturation and generation times limit their use and have precluded systematic genetic and mapping studies^5,21–23^. Here we describe a systematic cardiac screen of mouse lemurs, among the smallest, fastest reproducing, and most abundant non-human primates^5^, and show they are a tractable and valuable genetic model organism for cardiac arrhythmias.

Cardiac arrhythmias, irregularities of the heartbeat, afflict millions of people worldwide and cause one-fifth of all deaths, many from sudden cardiac death^1^. Arrhythmias result from alterations or disruption of the cardiac conduction system, which controls the regular and rhythmic contraction of the atrial and ventricular heart chambers that pump blood to the lungs and then on to the rest of the body to circulate oxygen, nutrients, and other small molecules and blood cells. As in other areas of medical research, mouse models have risen to prominence in arrhythmia research^2,3^, especially since the mapping of human familial arrhythmia genes and identification of many as cardiac ion channels and transporters (“channelopathies”) whose mouse homologs were then similarly altered by gene targeting^24,25^. But the substantial differences between mice and humans in conduction system structure, heart rate, action potential waveforms, and channel genes^26,27^, and the lack of naturally-occuring arrhythmias in mice and failure of some targeted mutations to mimic the human arrhythmia, have spurred the search for better and complementary models^4^. There is a pressing need for such models because despite the progress in elucidating conduction system genes, mechanisms and therapeutics, the pathophysiology of arrhythmias is typically complex and remains poorly understood and often difficult to treat^28^. Most available therapies are symptomatic so do not address the underlying defect, and unfortunately are ineffective in many patients. Although the rapidly improving ability to create, genetically modify, and interrogate human conduction system cell types in vitro holds great promise for the field (see below), it is not currently possible to recapitulate all the relevant cardiac cell types and other interacting cells that contribute to conduction system function and pathology in vivo.

Many aspects of mouse lemur physiology, behavior, ecology, and phylogeny have been elucidated over the past half century by studies in their native rainforests in Madagascar and in laboratory colonies in Europe and the United States^5^. To determine if mouse lemurs could be used for genetic interrogation of primate biology, behavior, and diseases, we and our collaborators systematically screened laboratory and wild mouse lemurs to find individual, heritable traits that could be genetically and molecularly mapped, the classical genetic approaches that have proven so powerful for the canonical genetic model organisms and for human diseases^9,10,29^. We phenotyped hundreds of individual lemurs using dozens of assays for diverse anatomical, physiological, and behavioral traits, like the phenotyping assays used to characterize targeted gene knockouts in mice (IMPC)^30^. Here we describe the results of an electrocardiograph (ECG) screen of lemurs to detect cardiac pathologies, and the subsequent pedigree analysis and genetic mapping of one of the eight identified familial, human-like arrhythmias. This uncovered a new gene, molecules, and mechanism of the primate cardiac pacemaker.

## RESULTS

### Mouse lemur cardiovascular phenotyping by electrocardiography

The cardiac conduction system of mammals has three major components (Fig. 1d)^31,32^. One is the sinoatrial node (SAN), electrically specialized cardiomyocytes that autonomously and rhythmically generate the action potentials that initiate each cardiac cycle (heartbeat). SAN cells serve as the pacemaker that normally dictates heart rate, increasing or decreasing their firing rate to match physiological demand. These electrical signals spread through and contract atrial cardiomyocytes, reaching the atrioventricular node (AVN) and His-Purkinje system (left and right bundle branches, LBB and RBB, and Purkinje fibers), the other two components that rapidly spread the contraction signal to ventricular cardiomyocytes to complete the cardiac cycle. Genetic, physical, or chemical alteration, dysregulation, or destruction of any of these three components can cause arrhythmias, as can creation of ectopic sites of cardiac electrical activity or repolarization^31^. The specific site and component affected, and how it is affected, determines the type and clinical significance of the arrhythmia, and its prognosis and therapies. Many arrhythmias and other cardiac pathologies can be detected by electrocardiography (ECG), which uses sensitive electrodes placed on the body surface to measure electrical changes in the heart, including depolarization and repolarization of the conduction cells and contracting cardiomyocytes as the pacemaker signal propagates through the heart during each beat.

**Figure 1.**
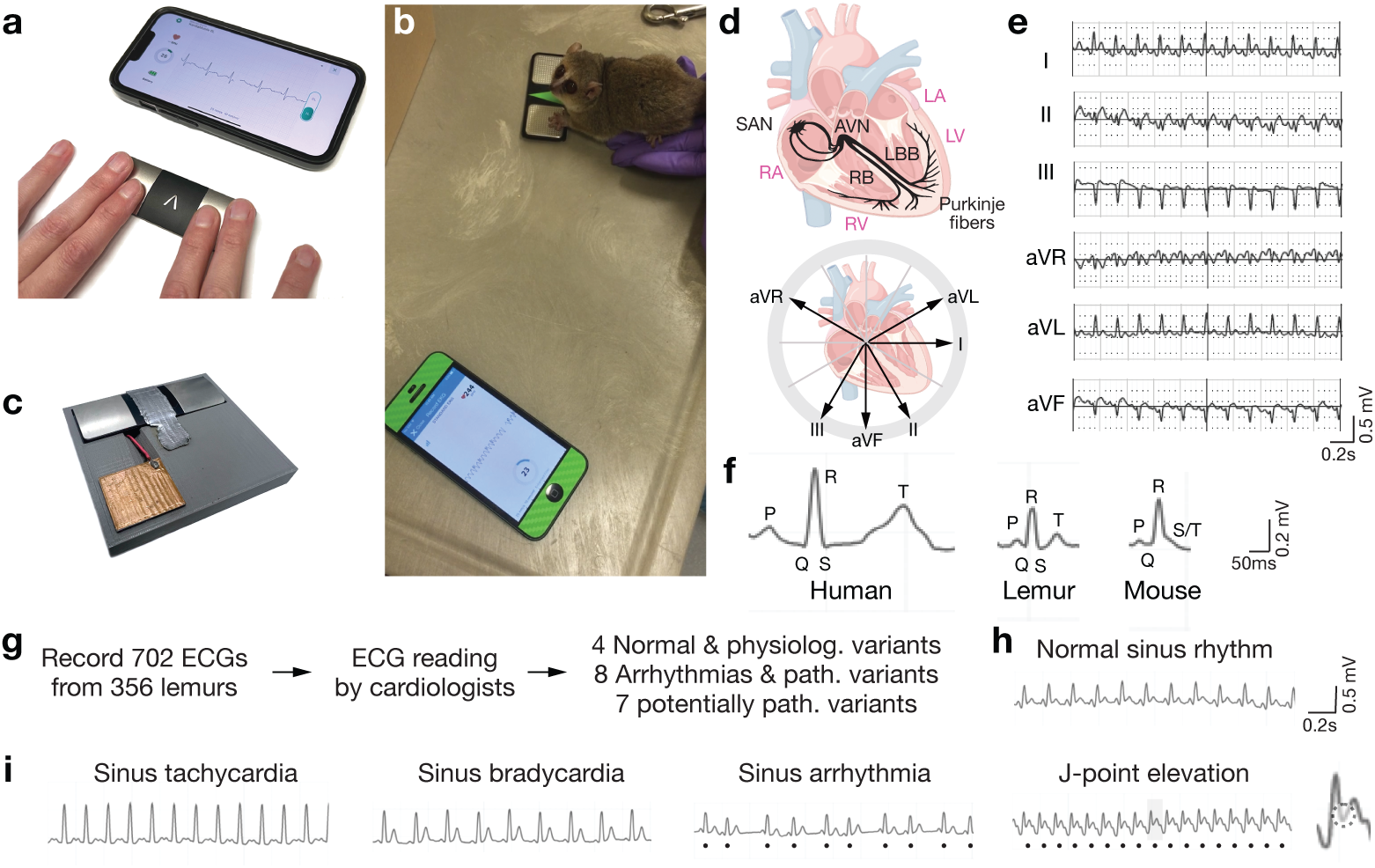
Systematic cardiovascular screening of mouse lemurs by point-of-care electrocardiography (ECG). **(a)** Portable human ECG monitor (KardiaMobile®, AliveCor) coupled to a smartphone app designed for ECG recording in real time at home. Device measures electric field differences between fingers placed on each conductive electrode pad that result from myocardial cell depolarization and repolarization spreading across the heart during each cardiac cycle (heartbeat). **(b)** Real time ECG recording of a resting (unanesthetized) mouse lemur as in panel **a** except the lemur’s hands contact the electrode pads. Image is a single frame taken from a live video recording (Movie S1). **(c)** Custom modified KardiaMobile 6L® ECG monitor with a third electrode pad (brown) on its surface. This enables 6-lead mouse lemur ECG detection (as shown in panel e) when the lemur’s foot contacts the third pad while its hands contact the other two pads as above. **(d)** (Top) Human cardiac conduction system (black). The sinoatrial node (SAN) pacemaker in the right atrium (RA) fires rhythmically to initiate each cardiac cycle, and this contraction signal spreads to the atrioventricular node (AVN) and through the His-Purkinje system (right and left bundle branches (RBB and LBB) and Purkinje fibers) to initiate contraction of the right and left ventricles (RV, LV). (Bottom) The six classical ECG limb leads^74^. Electric field changes caused by the spreading wave of myocardial depolarization and repolarization during the cardiac cycle are captured by the three electrode pads and integrated to derive the six electrical vectors shown: limb leads I, II, III, aVR, aVL, and aVF. **(e)** A 6-lead lemur ECG trace (2 seconds) captured by the customized device in panel **c**. Scale bar for ECG trace, 0.2 seconds, 0.5 millivolts. **(f)** Single cardiac ECG waveforms of human, lemur, and mouse captured with the six lead device. The three waves correspond to atrial depolarization (P), ventricular depolarization (QRS), and ventricular repolarization (T). Note the mouse lemur waveform resembles the human waveform, but in mouse there is no discrete T wave (fused with QRS). Scale bars for ECG traces, 50 milliseconds, 0.2 millivolts. **(g)** Scheme of the lemur ECG screen and overview of results. **(h)** ECG tracing (2 seconds) of lemur normal sinus rhythm. **(i)** ECG tracings of the four physiological variations detected: three heart rate variations (sinus tachycardia, sinus bradycardia, sinus arrhythmia) and J-point elevation, a typically benign waveform variant indicating early repolarization in younger individuals. Dots (sinus arrhythmia), R-waves of each heartbeat to highlight the beat-to-beat variability from rapid changes in vagal tone during inspiration and expiration. Dots (J-point elevation), elevated take-off of each ST-segment (see inset).

For rapid, point-of-care mouse lemur cardiovascular phenotyping, we used a simple commercial human home ECG monitor in which a mouse lemur subject was placed on the device with each hand positioned on an electrode pad (Fig. 1b), where fingers of a human subject are normally placed (Fig. 1a). The animals are quite docile, even wild mouse lemurs (see below), most needing only gentle guidance for hand placement, which typically remained in place for the full recording period (10 to 30 secs) (Movie S1). This allowed capture of mouse lemur electrocardiogram traces (Fig. 1e) in the subject’s native state, without need for sedation or anesthesia, cardiorespiratory-depressing agents that are typically required for ECG assessment of mouse and other animals^33,34^. We modified the standard commercial instrument by putting a third electrode pad near the others (Fig. 1c), on which a foot of the lemur was positioned. This provides a 3-electrode ECG (two hands and a foot), from which a conventional 6-lead ECG can be derived (Fig. 1d,e).

A typical cardiac cycle from an ECG of a laboratory *M. murinus* lemur displays the classical features of a healthy cardiac waveform in humans (Fig. 1f). Each cycle comprises three sequential electrical waves: the P-wave (representing atrial depolarization), QRS-complex (ventricular depolarization), and T-wave (ventricular repolarization). The *M. murinus* waveform resembles that of human more closely than mouse does: its T-waves are, like human, more curved and rounded, and there is a discrete T-wave separated from the QRS-complex as for human, whereas there is no discrete T-wave in mouse because ventricular repolarization begins immediately after the R-wave so the T-wave is partially fused with the QRS-complex (Fig. 1f). This suggests that chamber-specific ion channel expression and function may be better conserved between the two primates than between either primate and mouse. Other differences between mouse and human conduction systems are summarized in Table S1.

In the normal lemur heart at rest (normal sinus rhythm, NSR), each cardiac cycle was detected discretely from the preceding and successive beats. The heart rate (HR) for *M. murinus* was 431 ±61 beats per min (bpm, mean ±SD, n=334; range 309 - 553 (95% confidence interval), comparable to the value (427 ±43 bpm) for lab *M. murinus* obtained by continuous 1-lead invasive cardiac telemetry^35^. We determined NSR as 309 - 389 bpm (see Methods), with values outside this range demarcating slower (sinus bradycardia, SB) or faster (sinus tachycardia, ST) heart rates. Using our portable ECG monitor in and around Ranomafana Rainforest in Madagascar, we obtained slightly slower NSR values for wild *M. rufus* (260 - 458 bpm, 95% CI with mean 359 ±49, n=65) and a newly discovered but as yet unnamed species that we refer to here as *M. species nova* or *M. nova* for short (237 - 421 bpm, 95% CI with mean 329 ±46, n=37). The normal mouse lemur heart rate is intermediate between that of humans (60 - 100 bpm)^36,37^ and mice (600 - 800 bpm)^4^. The slower heart rate of mouse lemurs than mouse is valuable experimentally because it is less prone to artifacts and easier to detect arrhythmias and other ECG variants, and it may predispose to naturally-occurring arrhythmias (see below) that so far remain elusive in mice.

### ECG screening identifies eight lemur cardiac arrhythmias and pathological variants

The point-of-care ECG monitor was used to evaluate cardiac function in 356 mouse lemurs. We performed a total of 702 ECGs: 565 on 255 *M. murinus* lemurs from MNHN laboratory colony, and 137 ECGs on 101 wild *M. rufus* and *M. nova* lemurs (Fig. 1g). ECGs were typically repeated for each animal on the same day and, when possible, on separate days for corroboration; serial recordings were generally consistent except for gradually slowing heart rates indicating acclimation of the subject to the ECG procedure. Some lemurs were followed longitudinally, including ones with detected ECG pathologies (see below). Automated human ECG readers did not reliably interpret mouse lemur ECGs, so each ECG was independently read and validated by two to three board-certified cardiologists. This identified 19 different naturally-occurring mouse lemur arrhythmias and ECG variants (Fig. 1g), each resembling a known human condition including eight human arrhythmias and pathological variants (Fig. 2), seven other ECG variants that can be pathological in humans but require further diagnostic tests (Fig. 3), and four physiological or benign human variants (Fig. 1i).

**Figure 2.**
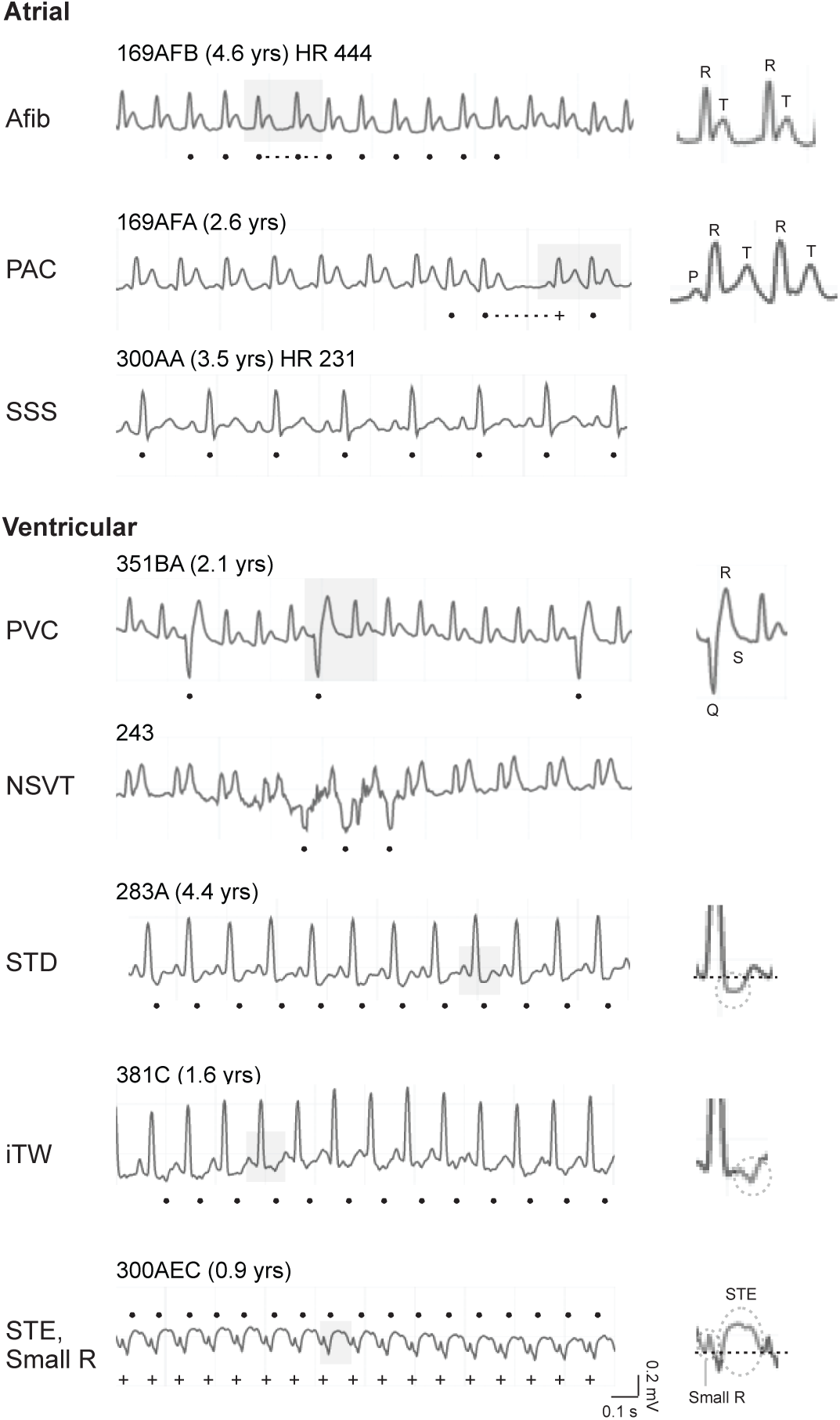
Eight cardiac arrhythmias and other pathologies identified in lemur ECG screen. ECG traces (2 seconds) and waveform pathologies (insets at right) of ECG variants detected in the screen that resemble ECG abnormalities of human arrhythmias and pathologies, the basis for the assigned names of the lemur ECG variants. The top three pathologies arise in the atria; the bottom five are ventricular pathologies. Lemur ID, age (if known) when ECG was taken, and heart rate (HR, in beats per minute, bpm, if integral to the arrhythmia diagnosis) are indicated above each trace. Dots, pathological beats. Gray shading, pathological beats magnified at right to highlight abnormality. Atrial fibrillation (Afib): Dots mark the R-wave of abnormal beats. Note the variable intervals between R-waves (e.g., dashed lines), illustrating the classic irregularly irregular rhythm of atrial fibrillation. Inset, most pronounced irregularity. Note lack of sinus P-waves, another hallmark of Afib. Premature atrial contraction (PAC): early beats (dots) that arise from an ectopic atrial focus and commonly result in a compensatory pause (dashed extension from dot) before the next normal beat (+) resumes. Inset, normal sinus beat followed by a PAC. Note absence of the ectopic P-wave of the latter, which is fused with the T-wave of the former. Sick sinus syndrome (SSS): an episodic, substantially reduced heart rate (bradycardia) that does not meet physiological need (see Fig. 4). Premature ventricular contraction (PVC): each PVC (dot) arises from an ectopic ventricular focus. Inset, classic wide and unusual shape of the QRS-complex, which is followed by a normal sinus beat. Non-sustained ventricular tachycardia (NSVT): the three consecutive PVCs (dots) represent an episode of NSVT. ST depression (STD): each beat shows ST-segment depression (dots), in which the initiation of the ST-segment is below the isoelectric point (dashed line in inset). Inverted T-wave (iTW): each beat shows an iTW (dots), in which T-wave orientation is flipped 180 degrees from normal (inset). ST elevation (STE): each beat shows ST-segment elevation (dots above trace), in which initiation of the ST-segment is above the isoelectric point (dashed line, inset). When STE is accompanied by a small R-wave (+ below trace), this can represent an evolving myocardial infarction (in humans). Scale bar for ECG traces, 0.1 seconds, 0.2 millivolts.

**Figure 3.**
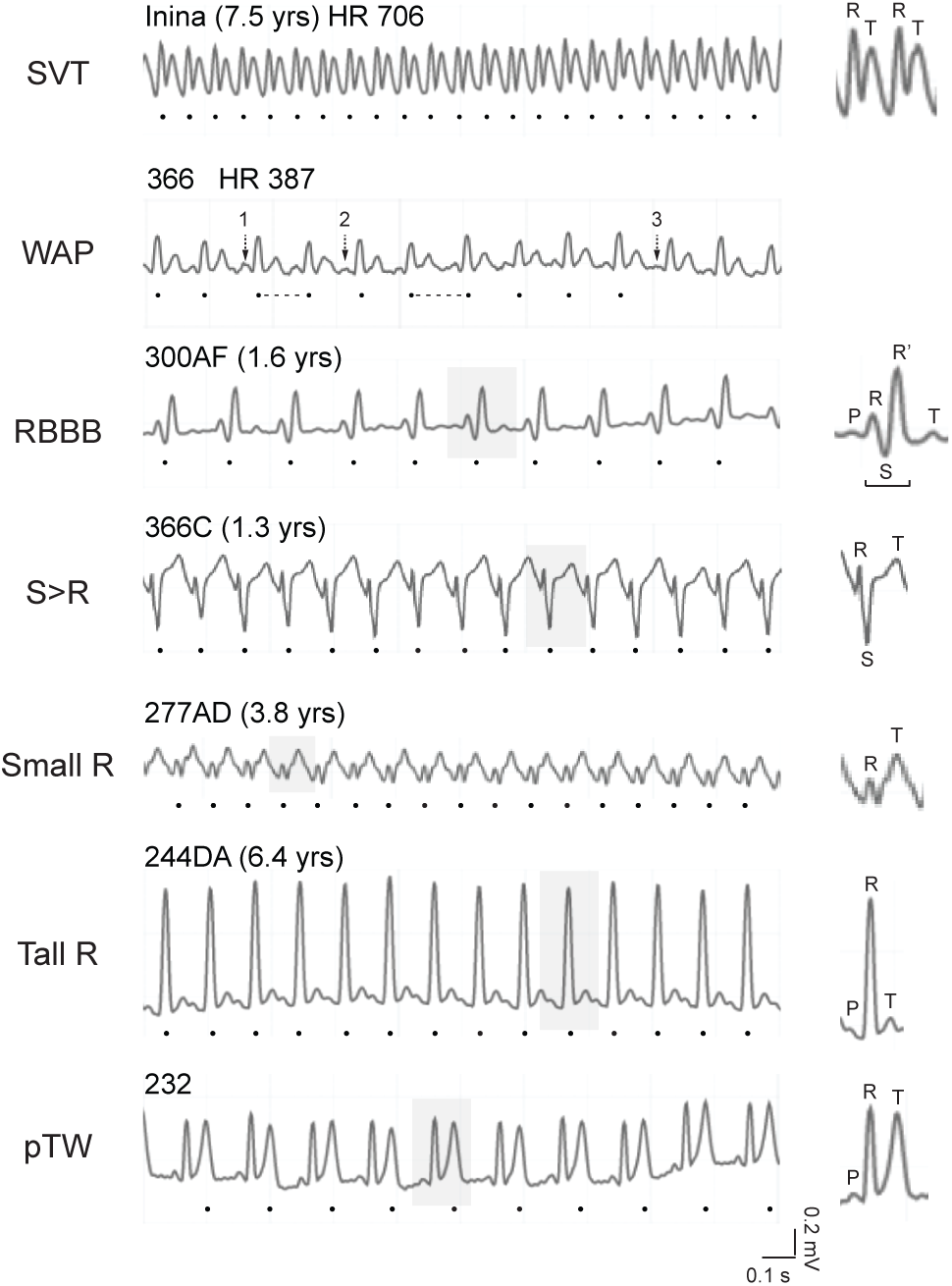
Seven potentially pathologic cardiac variants identified in the lemur ECG screen. ECG traces (2 secs) of lemur cardiac variants identified in the ECG screen as in Fig. 2 but resembling human variants that can be pathological but require additional clinical testing for diagnosis. Supraventricular tachycardia (SVT): a substantially increased heart rate with no P-waves detected (dots mark R-wave of abnormal beats). Wandering atrial pacemaker (WAP): an irregularly irregular rhythm (dots) in which different (“wandering”) ectopic atrial foci initiate beats. Note the variable R-R intervals (dashed lines show the longest intervals) and three distinct P-wave morphologies (numbered 1 - 3) resulting from the wandering foci. Right bundle branch block (RBBB): prolonged QRS-complex (dots) from delayed depolarization through the right bundle branch. Inset, single cardiac beat showing an abnormal R-S-R’ of a prolonged QRS complex (bracket). Large S and small R-waves (S>R): amplitude of S-wave (dots) is greater than that of R-wave. Isolated small R-waves (Small R): small R-wave amplitude (dots) relative to the T-wave. Tall R-waves (Tall R): tall R-wave (dots) relative to the P- and T-waves. peaked T-waves (pTWs): T-wave amplitude (dots) approaches that of the R-wave. Scale bar for ECG traces, 0.1 seconds, 0.2 mV.

Among the 356 mouse lemurs screened, 22 (6.2%) had ECG abnormalities resembling pathological human arrhythmias or variants (Table 1). Five lemurs (1.4%) had more than one ECG-diagnosed pathology including three lemurs with three different pathologies, indicating serious underlying cardiovascular disease (see below). The identified cardiac pathologies included major arrhythmias arising from both the atrial and ventricular chambers. One detected atrial arrhythmia was atrial fibrillation (Afib), identified in a 4.6 year old male (169AFB) in the lab colony (Fig. 2; typical mouse lemur lifespan ∼6 years^38^). Afib is the most common human arrhythmia^39^, marked by chaotic activation of multiple pathological atrial foci causing both atria to fibrillate and flow to stagnate, predisposing to blood clots and stroke. Hyperlipidemia-induced atherosclerosis, obesity, and premature atrial contractions (PACs), which arise in humans from extranodal atrial tissue that fires before and suppresses sinoatrial node pacemaking, are all significant Afib risk factors in humans^31^. The afflicted lemur (169AFB) showed several risk factors including PACs detected on ECG a year earlier (age 3.1). PACs were also found in two other lab lemurs and four wild lemurs (Table 1). A third atrial arrhythmia, resembling human sick sinus syndrome (SSS), was found in eight lemurs and is described in a separate section below.

**Table 1.** Mouse lemurs with cardiac arrhythmias and ECG pathologies. Deep phenotyping data of 22 animals (15 laboratory, 7 wild) with human-like ECG abnormalities. Lemurs were systematically screened (see Methods) including for the following parameters: family history, physical exam, body measure, behavior and strength assessments, serum lipids, and ECG. Dx, age in years of diagnosis; PAC, premature atrial contractions; Afib, atrial fibrillation; SB, sinus bradycardia; STD, ST-segment depression; iTW, inverted T-waves; PVC, premature ventricular contractions; NSVT, non-sustained ventricular tachycardia; STE, ST-segment elevation; Small R, small R-wave; Fam hx, family history; SSS, sick sinus syndrome (used interchangeably with tachy-brady syndrome).

| Name | Dx age (yr) | Sex | Lab vs wild | Pathology | Other clinical findings |
| --- | --- | --- | --- | --- | --- |
| 169AFA | 2.6 | Male | Lab | PAC | Fam hx of PAC/Afib |
| 169AFB | 4.6 | Male | Lab | Afib, PAC | Fam hx of PAC/Afib, hyperlipidemia, hyperglycemia |
| 262DBA | 3.5 | Male | Lab | PAC | Fam hx of SSS, low activity |
| 91 |  | Male | Wild *rufus* | PAC |  |
| 242 |  | Female | Wild *rufus* | PAC |  |
| 144 |  | Female | Wild *sp nov* | PAC |  |
| 317 |  | Female | Wild *sp nov* | PAC | Low activity |
| 204FD | 1.5 | Female | Lab | SB | Fam hx SSS, very low activity |
| 241EA | 4.6 | Female | Lab | SB (4.6), STD (4.6), iTW (4.6) | Fam hx SSS, low activity |
| 261BBA | 4.4 | Male | Lab | SB | Fam hx SSS, low activity, decreased grip strength |
| 300AA | 3.4 | Male | Lab | SB/tachy-brady | Fam hx SSS, very lethargic, very low activity, decreased grip strength |
| 300AB | 3.4 | Male | Lab | SB/tachy-brady (3.4), STD (3.5), iTW (3.5) | Fam hx SSS, very lethargic, sudden death during anesthesia |
| 300AC | 2.5 | Female | Lab | SB/tachy-brady (2.5), STD (3.4), iTW (2.5) | Fam hx SSS, very lethargic, low activity |
| 397C | 1.5 | Female | Lab | SB |  |
| 348 |  | Female | Wild *sp nov* | SB |  |
| 351BA | 2.1 | Male | Lab | PVC |  |
| 243 |  | Male | Wild *sp nov* | NSVT |  |
| 283A | 4.4 | Male | Lab | STD | Persistent tachycardia |
| 308A | 6.2 | Male | Lab | STD (6.2), iTW (6.2) | Persistent tachycardia, hyperglycemia |
| 308 |  | Male | Wild *sp nov* | STD |  |
| 381C | 1.6 | Female | Lab | iTW | Fam hx of SSS, persistent tachycardia, euthanized for weakness/lethargy |
| 300AEC | 0.9 | Female | Lab | STE (0.9), small R (0.9) | Tachycardia |

The screen also identified five arrhythmias and ECG pathologies of ventricular origin (Fig. 2, bottom). A 2.1 yr old lab lemur (351BA) showed premature ventricular contractions (PVCs), extra beats that arise from extranodal ventricular tissue in humans and form broad and bizarre QRS complexes like those seen in this lemur (Fig. 2). Such complexes in humans typically indicate cardiac pathology, notably myocyte irritability from ischemia or from structural alterations caused by cardiomyopathies, myocarditis, or valvular heart disease^31^. A PVC burden (fraction of normal beats) of 5% or more in human patients is considered high and places them at risk for sustained ventricular tachycardia (VT) and arrhythmia-induced pathological ventricular remodeling and heart failure^40^; the PVC burden of the afflicted lemur (351BA) was 23%, over four times this human clinical threshold. A wild *M. nova* male (243) had non-sustained ventricular tachycardia (NSVT), three or more consecutive PVCs (Fig. 2). This pathology in humans most commonly results from coronary ischemia, and if sustained (VT), leads to decreased cardiac output, hypotension, and syncope; VT is the leading cause of sudden cardiac death in adults^41^.

ST-segment depression (STD) and inverted T-waves (iTW) are ventricular repolarization pathologies that in human patients commonly arise from myocardial ischemia from coronary artery disease. Four lab lemurs showed both of these ECG pathologies. Three of them were also afflicted with the episodic bradycardia of sick sinus syndrome (see below), whereas 308A was persistently tachycardic (Table 1). One lab lemur (283A, Fig. 2) and one wild lemur (308, Table 1) showed STD alone, and another lab lemur (381C, Fig. 2) showed only iTW, detected when she was just 1.6 yrs old. 283A was persistently tachycardic on all other ECGs (3 on 3 different days, avg 463 bpm) and 381C was tachycardic on all ECGs (4 on 4 different days, avg 458 bpm). One young lab lemur (300AEC) was tachycardic (500 bpm) with ST-segment elevation (STE) in conjunction with a small R-wave (Fig. 2; see below), which in humans diagnosed on 12-lead ECG is highly sensitive for acute myocardial infarction, a cardiac emergency known as ST-elevation myocardial infarction (STEMI) where the infarct spans the full thickness of the heart wall (transmural)^31^. STEMI manifests classically with elevation of the ST-segment in a convex manner (resembling a tombstone), and a small R-wave due to loss of R-wave voltage as the myocardium becomes necrotic, both seen here in lemur 300AEC’s ECG.

### Identification of seven potentially pathological lemur ECG variants

In addition to the arrhythmias and pathological variants described above, we identified four physiological or benign variants in heart rate and waveform (Fig. 1i) along with seven potentially pathological variants that can be pathological in humans but depend on other physical symptoms and signs for definitive diagnosis (Fig. 3).

Potentially pathological variants include supraventricular tachycardia (SVT), found in 14 lab lemurs. In humans, SVT can result either from high but benign activity of the sinoatrial node pacemaker that obscures the normal P-wave, or from the pathological activity of extranodal atrial tissue driving an atrial tachyarrhythmia such as atrial fibrillation. Another potentially pathological variant we uncovered was wandering atrial pacemaker (WAP). This is usually a benign condition in humans found incidentally in young, healthy athletic individuals caused by changes in vagal tone (e.g., from stress, diet, transient electrolyte disturbances) that stimulate ectopic foci and produce an irregularly irregular rhythm^31^. However, it can also occur in the elderly caused by sinus node dysfunction or chronic lung conditions. Two wild lemurs were identified with WAP, and both also had intermittent peaked T-waves (see below), which too is often a normal variant in young patients or caused by electrolyte changes. One lab lemur (300AF, 1.6 yrs old) was found with right bundle branch block (RBBB), a widening of the QRS complex caused by delayed depolarization of the right bundle branch of the conduction system. Although RBBB in humans is typically benign^42^, it is a common presentation of several congenital heart diseases and can be a sequela of several primary cardiac and lung diseases. Large S and small R-waves (S>R) are associated with right deviation of the cardiac axis in humans from right heart strain, and was found in four lab lemurs. Small R-waves were found in 17 lab lemurs and one wild lemur. This is a common ECG finding in humans resulting from loss of cardiac tissue from myocardial infarction or cardiac fibrosis, or from normal variation in heart orientation^31^. Three lab animals (300AEC, 300AED, 300AEB) had small R-waves at young ages. For 300AEC, this co-presented with STE (see above), while small R-wave alone was seen in 300AEB and 300AED (at ages 0.9 and 3.0, respectively). Tall R-waves were found in 32 lab lemurs and 6 wild lemurs. In humans this is commonly caused by thickening of the myocardium from cardiomyopathy or pathological hypertrophy, but it can also be seen with performance hypertrophy (“athlete’s heart”) or thin chest wall. Peaked T-waves (pTWs) were observed in 27 lab lemurs and 40 wild lemurs. In humans this can be a normal variant in youth and athletes^43^, but it can also result from acute myocardial infarction or hyperkalemia, the latter potentially contributing to its surprising prevalence in wild lemurs that are baited with a high potassium (banana) meal.

Table 2 compares the prevalence of cardiac arrhythmias and variants in laboratory and wild mouse lemurs. There were no statistically significant differences in prevalence of any of the rare arrhythmias or pathological variants, or the overall prevalence of cardiac arrhythmias and pathological variants between lab (5.9%, 15 of 255 screened) and wild (6.9%, 7 of 101) lemurs. Although a similar unbiased, systematic ECG screen has not been done for humans, the prevalence of human arrhythmias is estimated at 2-5% of the population^39^, similar to the value obtained here for lemurs. Curiously, in the lemurs there were striking prevalence differences in one common potentially pathological variant (pTWs: 11% lab vs 40% wild, p<1.6e-3) and one normal variant (sinus arrhythmia, SA: 2.0% vs 43%, p<1.6e-3), perhaps because young lemurs are overrepresented among wild lemurs and these variants are more prevalent in young lemurs as they are in young humans. There were no statistically significant differences in prevalence between sexes in either lab or wild animals.

**Table 2.** Prevalence of mouse lemur arrhythmias and ECG variants in wild and laboratory mouse lemurs. Values shown are number (n) and percent individuals affected for each species and screening site (and totals): *M. murinus* lab mouse lemurs (MNHN, Brunoy, France; 255 screened), *M. rufus* and *M. nova* wild mouse lemurs (Ranomafana National Park, Madagascar; 101 mouse lemurs screened). Values are stratified by sex (F, female, M, male). Overall prevalence was 8.4% (30 diagnosed arrhythmias), which were found in 6.2% of all mouse lemurs screened (22 mouse lemurs, 5 showed >1 ECG abnormality). PACs, premature atrial contractions; Afib, atrial fibrillation; SSS, sick sinus syndrome; PVCs, premature ventricular contractions; NSVT, non-sustained ventricular tachycardia; STD, ST-segment depression; iTWs, inverted T-waves; STE, ST-segment elevation; SVT, supraventricular tachycardia due to an unknown underlying rhythm; WAP, wandering atrial pacemaker; RBBB, right bundle branch block; S>R, S wave larger than R wave; Small R, small R-wave; Tall R, tall R-wave; pTWs, peaked T-waves; L-voltage, low voltage.

| Arrhythmias<br>& ECG pathologies | Lab |  |  | Wild |  |  |  |  |  |  | Lab +<br>Wild |
| --- | --- | --- | --- | --- | --- | --- | --- | --- | --- | --- | --- |
|  | *M. murinus* |  |  | *M. rufus* |  |  | *M. nova* |  |  |  | (356) |
|  | F<br>(136) | M<br>(119) | F+M<br>(255) | F<br>(33) | M<br>(31) | F+M<br>(64) | F<br>(19) | M<br>(18) | F+M<br>(37) | All<br>wild<br>(101) |  |
|  | % (n) | % (n) | % | % (n) | % (n) | % | % (n) | % (n) | % | % | % |
| PACs | 0 | 2.5 (3) | 1.2 | 3.0 (1) | 3.2 (1) | 3.1 | 11 (2) | 0 | 5.4 | 4.0 | 2.0 |
| Afib | 0 | 0.8 (1) | 0.4 | 0 | 0 | 0.0 | 0 | 0 | 0.0 | 0.0 | 0.3 |
| SSS | 2.9 (4) | 2.5 (3) | 2.7 | 0 | 0 | 0.0 | 5.3 (1) | 0 | 0.0 | 1.0 | 2.2 |
| PVCs | 0 | 0.8 (1) | 0.4 | 0 | 0 | 0.0 | 0 | 0 | 0.0 | 0.0 | 0.3 |
| NSVT | 0 | 0 | 0.0 | 0 | 0 | 0.0 | 0 | 5.6 (1) | 2.7 | 1.0 | 0.3 |
| STD | 1.5 (2) | 2.5 (3) | 2.0 | 0 | 0 | 0.0 | 0 | 5.6 (1) | 2.7 | 1.0 | 1.7 |
| iTWs | 2.2 (3) | 1.7 (2) | 2.0 | 0 | 0 | 0.0 | 0 | 0 | 0.0 | 0.0 | 1.4 |
| STE | 0.7 (1) | 0 | 0.4 | 0 | 0 | 0.0 | 0 | 0 | 0.0 | 0.0 | 0.3 |
| **TOTAL** | **7.4 (10)** | **11 (13)** | **9.0** | **3.0 (1)** | **3.2 (1)** | **3.1** | **16 (3)** | **11 (2)** | **14** | **6.9** | **8.4** |
| **ECG variants** |  |  |  |  |  |  |  |  |  |  |  |
| SVT | 7.4 (10) | 3.4 (4) | 5.5 | 0 | 0 | 0.0 | 0 | 0 | 0.0 | 0.0 | 3.9 |
| WAP | 0 | 0 | 0.8 | 0 | 3.2 (1) | 1.6 | 0 | 5.6 (1) | 2.7 | 2.0 | 0.6 |
| RBBB | 0.7 (1) | 0 | 0.4 | 0 | 0 | 0.0 | 0 | 0 | 0.0 | 0.0 | 0.3 |
| S>R | 2.9 (4) | 0 | 1.6 | 0 | 0 | 0.0 | 0 | 0 | 0.0 | 0.0 | 1.1 |
| Small R | 7.4 (10) | 5.9 (7) | 6.7 | 3.0 (1) | 0 | 1.6 | 0 | 0 | 0.0 | 1.0 | 5.1 |
| Tall R | 9.6 (13) | 16 (19) | 12.5 | 3.0 (1) | 13 (4) | 7.8 | 5.3 (1) | 0 | 2.7 | 5.9 | 11 |
| pTWs | 6.6 (9) | 15 (18) | 10.6 | 45 (15) | 32 (10) | 39.1 | 42 (8) | 39 (7) | 40.5 | 39.6 | 19 |
| SA | 0.7 (1) | 3.4 (4) | 2 | 61 (20) | 48 (15) | 54.7 | 32 (6) | 11 (2) | 21.6 | 42.6 | 13 |

Arrhythmias/pathologies and variants were detected using an adapted 6-lead human ECG assay (Kardia, AliveCor®, see Methods). This enabled successful diagnosis of the following arrhythmias (PACs, Afib, SSS, PVCs, NSVT, SVT, WAP, SA). Other diagnoses were made provisionally (STD, iTWs, STE, RBBB, S>R, Small R, Tall R, pTWs) with the 6-lead assay, but require further corroboration with standard 12-lead ECG. Prevalence comparisons of cardiac arrhythmias and variants between lemur sexes and species were determined by two-tailed Fisher's exact test. P-values were corrected for multiple hypotheses using the Holm-Šidák method. □
Abbreviations: ECG, electrocardiogram; F, female; M, male; (), number of female/male animals; % (n), % of animals with a given arrhythmia/pathology or variant, n = # of animals with a given arrhythmia/pathology or variant; PACs, premature atrial contractions; Afib, atrial fibrillation; SSS, sick sinus syndrome; PVCs, premature ventricular contractions; NSVT, non-sustained ventricular tachycardia; STD, ST-segment depression; iTWs, inverted T-waves; STE, ST-segment elevation; SVT, supraventricular tachycardia; WAP, wandering atrial pacemaker; S>R, large S- and small R-waves; Small R, small R-wave; Tall R, tall R-wave; pTWs, peaked T-waves; SA, sinus arrhythmia

### Lemur sick sinus syndrome

An atrial arrhythmia resembling human sick sinus syndrome (SSS) was identified in seven lab lemurs and one wild lemur (Fig. 2, Table 1). Human SSS is an episodic, substantially slow heart rate (bradycardia) that manifests symptomatically, affecting ∼1 in 600 adults over age 65^44,45^. Most commonly it results from scarring, degeneration, or damage to the sinoatrial node from aging or primary cardiac diseases such as atrial fibrillation, myocardial infarction, and cardiomyopathies, but there are also rarer genetic causes (see below). Pathological remodeling of the sinoatrial node over time can be complicated by overcompensatory activation of ectopic atrial foci, resulting in abnormal tachycardias, so SSS can present as (and is sometimes called) tachycardia-bradycardia syndrome (TBS) that alternates between periods of unusually slow and fast heart rates^45^. As the disease progresses, it can cause sinus arrest, lethargy, dizziness, confusion, and life-threatening syncope. It is typically treated by implantation of an electronic pacemaker device that rhythmically stimulates the heart to restore normal heart rate and function^45^.

In the seven affected lab lemurs, we found episodic, substantially slow heart rates that ranged from 169 - 300 bpm during the bradycardic episodes (Fig. 4, top). Between episodes, heart rates sometimes returned to normal, but three lemurs (300AA, 300AB, 300AC) showed tachycardia during an interim period as in human tachycardia-bradycardia syndrome. It was an early onset disease that was detected at late juvenile stage or early adulthood (age 1.5 – 4.6 yrs) (Table 1).

**Figure 4.**
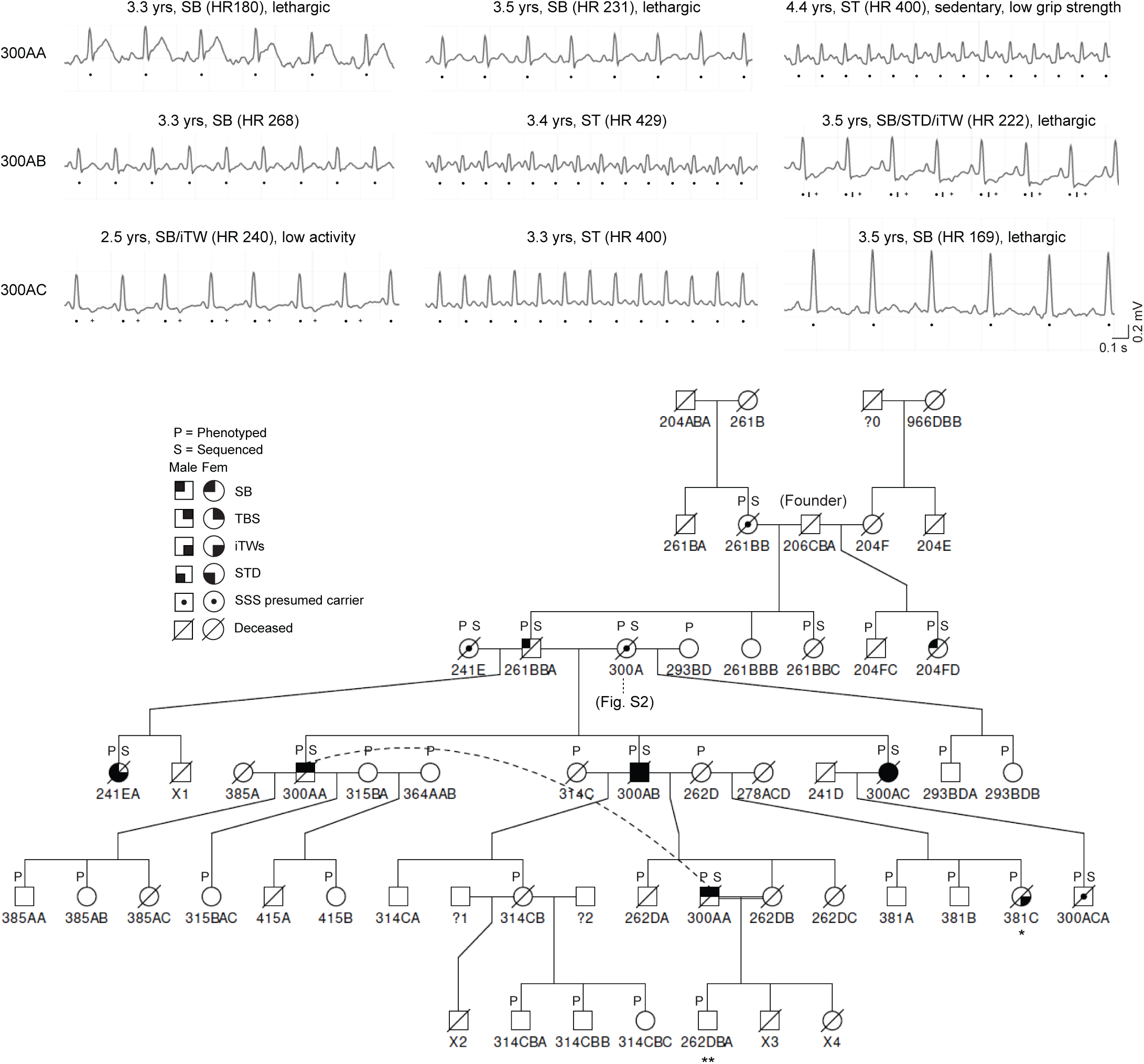
Lemur sick sinus syndrome pedigree. (Top) Longitudinal ECG studies (2 sec tracings) of three lemurs (name/ID number at left) with symptomatic sinus bradycardia (SB). Above each trace is the age when the ECG was taken, diagnosed ECG abnormalities and associated pathologies (SB, sinus bradycardia; ST, sinus tachycardia; STD, ST-segment depression; iTW, inverted T-wave), heart rate (HR, in bpm), and abnormal signs on physical exam. All three lemurs have tachy-brady syndrome (TBS), in which all ECGs showed either SB or ST (and none showed normal sinus rhythm). Dots, individual heartbeats during SB or ST. 300AB and 300AC manifest intermittently with a pathological ECG triad of SB (dots), STD (tick, not indicated for 300AC), and iTW (+). Scale bar, 0.1 seconds, 0.2 mV. (Bottom) Six generation pedigree of a lemur family that includes five individuals (the above three plus 261BBA and 241EA) with symptomatic SB and the associated pathologies or symptoms indicated by black fill of the individual’s symbol (see legend). Presumed carriers (inferred from the pedigree based on autosomal recessive inheritance and supported by genotyping in Fig. 5) are indicated by a central dot in the symbol. ?0, ?1, ?2, identity of father not yet resolved. Four lemurs were found dead in their cages at a young age: 261BA (3.6 yrs), 385A (2.9 yrs), 314C (2.2 yrs), 262DC (2.3 yrs). X1, X2, X3, X4, lemurs that died several days after birth (before naming). *, 381C diagnosed with iTW (age 1.6), persistent tachycardia, weakness/lethargy. **, 262DBA diagnosed with PACs and low activity (age 3.5).

Several associated clinical and ECG findings in the affected individuals suggest that the episodic bradycardia was clinically significant. First, all but one of the affected lemurs exhibited severe fatigue on physical exam or a combination of low activity scores and/or decreased grip strength in our deep phenotyping assay (Table 1), which commonly manifested at the same time or day as the recorded ECG pathology. Lethargy is often the presenting symptom of human SSS due to inadequate cerebral perfusion, warranting electronic pacemaker placement. Second, three of the affected individuals (241EA, 300AB, 300AC) showed additional ECG pathologies beyond the episodic bradycardia, notably ST-segment depression (STD) and inverted T-waves (iTW) (Fig. 4, Table 1). Both of these ECG pathologies signify abnormalities in ventricular repolarization that are commonly seen in states of myocardial ischemia in humans^31^, suggesting that the slow heart rate in these lemurs was not meeting cardiac demand. Consistent with this interpretation, the pathological ECG triad (SSS/STD/iTW) typically presented together on the same ECG traces (Fig. 4, top, 300AB age 3.5 trace). Finally, one of the affected lemurs (300AB, with the pathological triad), died prematurely at age 3.9 yrs. His sudden death during routine anesthesia suggests that the anesthetic (isoflurane, a cardiorespiratory depressant) may have caused cardiac arrest by further compromising his SSS chronotropic incompetence. Two other affected lemurs were found dead in their cages, and 314C, although not formally diagnosed with SSS because it was only assayed by ECG once at age 1.5 yrs, had borderline bradycardia (310 bpm) and grip strength, and was found dead at 2.2 yrs. Seven other animals in the SSS pedigree (see below, Fig. 4) also died young of unexplained death.

### Lemur SSS shows familial clustering

Lemur SSS was an early onset disease similarly affecting both sexes (Table 1), and the available serum profiles of affected individuals (n=4, 204FD, 241EA, 261BBA, 300AC) showed normal glucose levels and lipid panels including cholesterol. This suggested that advanced age and other common cardiovascular risk factors (hyperlipidemia, atherosclerosis) for the most common etiologies of human SSS did not underlie this lemur disease. The human disorder can also be a channelopathy caused by mutations in cardiac conduction channels HCN4^46,47^ or SCN5A^48^, which result in severe symptomatic bradycardia requiring pacemaker placement. To explore potential genetic causes of lemur SSS, we constructed a pedigree of the Brunoy laboratory colony (Karanewsky, Pendleton, Anzeraey, Terrien et al, unpublished data) and used it to determine if there was a family relationship among affected individuals. Maternal identities were ascertained from historical breeding records, but because paternity was ambiguous (for routine breeding, estrus females are typically presented with three or four male suitors) it was resolved by DNA sequencing of five selected polymorphic loci of each individual in the colony, its mother, and all possible fathers (see Methods).

The pedigree revealed striking familial clustering of lemur SSS (Fig. 4, bottom). Five of the affected lemurs are first degree relatives: an affected father (261BBA) and his two sons (300AA, 300AB) and daughter (300AC) from breeding with an unaffected mother (300A), as well as the daughter (241EA) of the same father (and half-sister of the others) from breeding with another unaffected mother (241E). The father had SB alone (mild phenotype), three children (300AA, 30300AB, 300C) had the progressively worse tachy-brady syndrome (TBS) of which two (300AB, 300AC) along with their half-sister (241EA) had the pathological triad (SB, iTW, STD). Determining the pedigree of a sixth affected individual (204FD) is in progress, but initial genotyping data indicate that he and 261BBA are likely half brothers, sharing the same father (206CBA); their presumptive father (206CBA) would then be the likely founder of this SSS pedigree. Pedigree construction is also underway for the seventh affected individual (397C), a potential two-generation descendant of the 300-designated animals (Fig. S1). Although thus far the next generation has not shown SSS on ECG screening, iTWs were detected at age 1.6 yrs in a female offspring (381C) of 300AB, who was persistently tachycardic and euthanized at 5.2 yrs due to weakness and lethargy. Two other offspring of his died prematurely of unknown causes: 262DC (2.3 yrs, before ECG screening was done) and 262DA (5.1 yrs), the latter with decreased grip strength (at 4.5 yrs).

We have also begun investigating family relationships of lemurs afflicted with other arrhythmias and CVDs, and uncovered a second ECG pathology with familial clustering. Two of the three lab lemurs identified in the screen with premature atrial contractions (PAC prevalence in colony, 2.5%), one of whom also progressed to atrial fibrillation, are littermates (169AFA, 169AFB) (Fig. S2). Like SSS, PACs (potentially progressing to atrial fibrillation) is thus also likely to be familial, possibly another simple Mendelian disease segregating in the lab colony. Other potentially pathologic ECG traits as well as a serum lipid disorder resembling human familial hypercholesterolemia uncovered in the deep phenotyping screen also showed familial clustering and will be described elsewhere.

### Lemur SSS is an autosomal recessive disorder that maps to a locus on chromosome 7 including *SLC41A2* magnesium transporter

The SSS pedigree suggested that this arrhythmia is a simple Mendelian disorder, consistent with either autosomal recessive or dominant inheritance pattern. To determine the inheritance pattern and map the SSS disease-causing locus, we performed Illumina paired-end, short-read shotgun whole genome sequencing (27x - 293x coverage, median 54x) for 35 lemurs from the lab colony including six lemurs with SSS (all except 397C) and 29 unaffected individuals (five unaffected family members plus 24 control animals outside the SSS pedigree) (Table S2). We used the obtained sequences to create an extensive library of *M. murinus* genomic sequence variants (∼45 million variants, one every ∼55 nucleotides on average), including mostly single-nucleotide polymorphisms (SNPs, 80%) but also short (typically <50 base pair) insertion (∼10%) and deletion (∼10%) alleles (Chang et al, unpublished data) that could be used for mutation mapping by linkage analysis (Fig. 5a).

**Figure 5.**
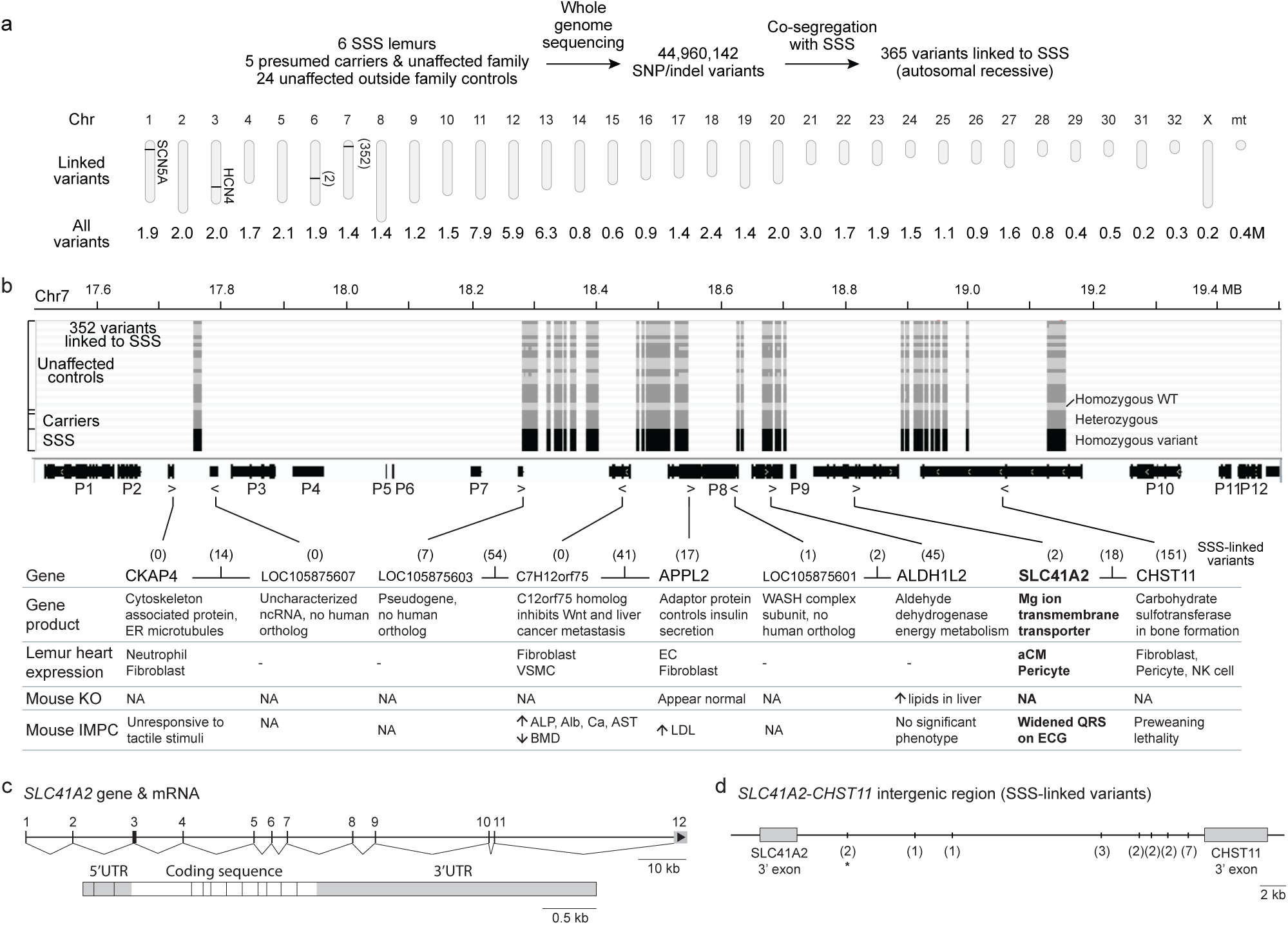
Genetic mapping of lemur sick sinus syndrome and the candidate genes in the mapped interval. **(a)** Genetic mapping scheme. Whole genome sequencing was done on: 6 of the 7 SSS lemurs identified in the screen, 5 presumed carriers and unaffected family members, and 24 unaffected (outside immediate family) controls in the lab colony (Table S2). Single nucleotide sequence polymorphisms (SNP) and insertion/deletion (indel) polymorphisms (variants) were identified across the genome (∼45M total), and those that co-segregated with the SSS trait based on autosomal recessive inheritance were identified (365 variants). The positions of 354 linked variants are shown on the mouse lemur diploid genome (32 autosomes, 1 pair of sex chromosomes, and a haploid mitochondrial genome); the remaining 11 variants mapped to two unplaced scaffolds (Table S3). The total number of variants identified on each chromosome (in millions, M), regardless of SSS linkage, is shown below each chromosome. The map locations of the lemur orthologues of the two genes (*SCN5A, HCN4*) known to cause human SSS are indicated. **(b)** Expanded view and graphic (Integrative Genomics Viewer) showing the variant genotypes of the 35 sequenced lemurs (rows) in the 1.4 Mb genomic interval on chromosome 7 containing the 352 variants linked to SSS (shaded columns). (Many variants are tightly clustered along the chromosome so at this resolution appear as a shaded block rather than a shaded thin line.) Each row in the plot shows the genotype (homozygous wild-type reference, light gray shading; heterozygous wild-type/variant, dark gray; homozygous variant, black) of one of the 35 sequenced lemurs whose SSS phenotype/carrier status is indicated at left. Note that all the SSS lemurs are homozygous variant (black fill) in this interval and all the carriers are heterozygous, whereas the unaffected controls are either homozygous wild-type reference or heterozygous, as expected for an autosomal recessive trait. The locations of the 21 genes (horizontal black bars) in the interval are shown below the graphic; thicker portions of the bars indicate exons, and arrowheads below bar show direction of transcription. The 9 candidate genes associated with or neighboring 1 or more of the 352 SSS-linked variants are named and described in the table below along with the number of associated variants (shown in parentheses at top of the table along with the number of intergenic variants). The other 12 genes (P1 to P12) with no associated or neighboring intergenic variants are similarly detailed in Table S4. The gene descriptions in the table include (top to bottom): gene product function (Gene product), lemur heart cell types in which the gene is highly and differentially expressed (Lemur heart expression) from the lemur transcriptomic atlas^49^ (“-” indicates gene is not detected or differentially expressed), and mouse knockout phenotypes of the orthologous gene in published literature (Mouse KO) or determined by the International Mouse Phenotyping Consortium (Mouse IMPC). **(c)** Structures of lemur *SLC41A2* gene (top) and spliced mRNA (bottom). Numbers, exons 1-11. White/no shading, protein coding sequence in mRNA; gray shading, non-coding sequences (5’-untranslated region, 5’UTR; 3’ untranslated region, 3’UTR). **(d)** Map of the 33,108 bp intergenic region between 3’ end of *SLC41A2* and 3’ end of *CHST11* showing positions of the 20 SSS-linked variants in this region. Numbers (in parentheses) indicate the number of linked variants clustered at that position; the 2 variants downstream and associated with *SLC41A2* (as assigned by SnpEff) are indicated by an asterisk; the other 18 are assigned as intergenic variants. Of these 20 variants, 10 are conserved between lemur and human including one of the *SLC41A2*-associated variants (Fig. S4).

We searched among the identified variants for ones that were tightly linked to (co-inherited with) the SSS phenotype in the pedigree (Fig. 5a). No variants were co-inherited with SSS in an autosomal dominant (AD) pattern, indicating the disease is unlikely to be a simple Mendelian autosomal dominant disorder. In contrast, we found 365 variants that were perfectly linked to the SSS phenotype in an autosomal recessive inheritance pattern (Table S3). Nearly all of the linked variants (352, >96%) mapped to a single genomic interval spanning 1.4 Mb on chromosome 7 (Fig. 5a,b); the rest (13) were scattered in several locations around the genome, all but two on short, unplaced scaffolds (Fig. 5a, Table S3). The mapped 1.4 Mb region contains nine candidate genes with associated variants, with half of the linked variants (171, 49%) localized to the ∼260 kb subregion at the right end of the interval containing two genes, *SLC41A2* and *CHST11*. None of the linked variants were in coding exons, except for a single synonymous variant (c.2157A>G, ALA719ALA) in *ALDH1L2* (Fig. 5b). This suggests that the disease-causing mutation is likely a regulatory mutation.

This linkage analysis and mapping of the lemur SSS locus excludes *HCN4* and *SCN5A*, orthologs of the two canonical human SSS channelopathy genes^46–48^, which in lemur are located on chromosomes 3 and 1 (Fig. 5a). Among genes within the mapped interval on chromosome 7 (Fig. 5b), the most appealing candidate was *SLC41A2*, a little studied transmembrane magnesium transporter^6,7^. It is the only gene in the interval that encodes an ion transporter or channel, and it is expressed in lemur atrial cardiomyocytes where the sinoatrial node pacemaker resides (Fig. 5b)^49^. A mouse *SLC41A2* knockout generated and systematically phenotyped by the Knockout Mouse Project (KOMP2) identified a cardiac ventricular conduction defect in the homozygous mutants that manifests as a widened QRS interval on ECG^8^ (Fig. S3). The phenotype is selective for heart and this specific conduction defect (Fig. S3), much as the lemur phenotype is selective for heart and the SSS arrhythmia and its sequelae. However, there was no evidence of a widened QRS interval in affected lemurs (32.6 ±1.2 msec, mean ±SD in n=7 affected lemurs vs. 32.4 ±1.8 msec in n=10 wild-type; p=0.87 (non-significant) by two-tailed student’s t-test with equal variance), and conversely there was no evidence of SSS or other arrhythmias or sinus node pacemaker defects in the mouse *SLC41A2* knockout (Fig. S3). Hence, if *SLC41A2* is the lemur SSS disease gene, then it functions differently in the mouse and lemur hearts. None of the other genes in the mapped lemur SSS disease interval had any obvious connection to cardiac pacemaking, conduction or disease, nor did any of the available mouse knockouts in these genes display a cardiac phenotype (Fig. 5b, Table S4).

### *SLC41A2* knockout slows firing and alters magnesium dynamics of human SA node pacemaker cells

To determine if *SLC41A2* functions in the cardiac pacemaker of primates, we developed the scheme shown in Figs. 6c and S6a,b to create a CRISPR/Cas9-mediated knockout of *SLC41A2* in SA node cells (iSANCs) derived from human pluripotent cells. DNA sequencing of the obtained knockout iSANCs showed that mutant cells had a 118 bp homozygous deletion in the targeted exon (exon 2) of *SLC41A2*, resulting in a frameshift mutation and early stop codon (Figs. 6a, S6b). Immunostaining detected expression of SLC41A2 in wild-type iSANCs (see below) but not in the *SLC41A2* knockout iSANCs (Fig. S6c).

**Figure 6.**
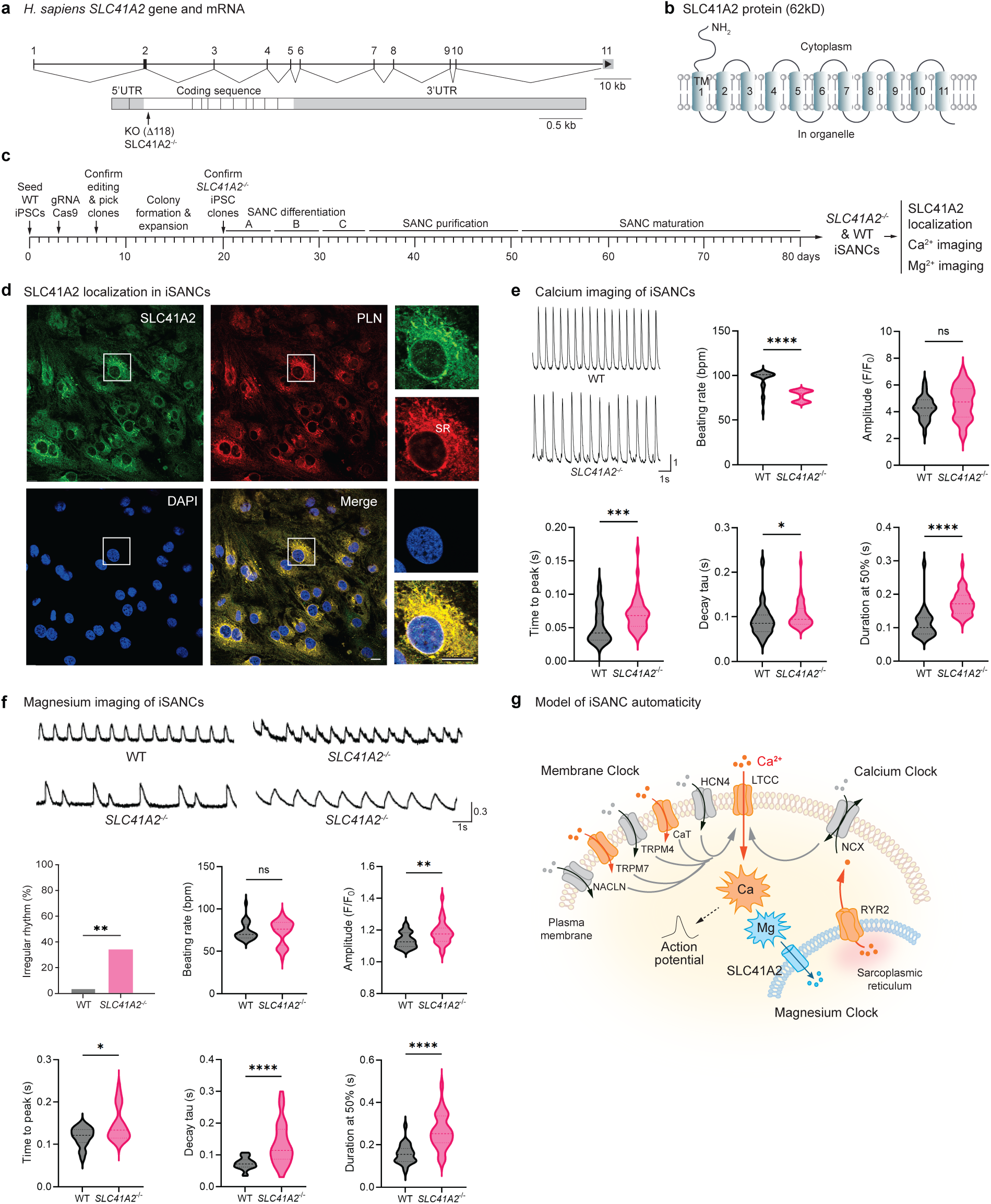
Expression and knockout of *SLC41A2* in human sinoatrial node cells. **(a)** Structures of human *SLC41A2* gene (top) and spliced mRNA (bottom) and the site of the introduced *SLC41A2* gene knockout (KO) in human iPSC-derived sinoatrial node cells (iSANCs). The *SLC41A2* KO (SLC41A2^-/-^) is an 118 bp deletion in exon 2, causing an early truncation and frameshift of the coding sequence (Fig. S6b), which was created in human induced pluripotent stem cells (iPSCs) by CRISPR-Cas9-mediated gene targeting prior to their differentiation into iSANCs, as described in panel c and Figure S6a,b. Numbers, exons 1-10. Dark blue shading, protein coding sequence in mRNA; light blue shading, non-coding sequences (5’-untranslated region, 5’UTR; 3’ untranslated region, 3’UTR). **(b)** Model of the membrane topology of human SLC41A2 with 11 transmembrane (TM) domains, modified from Sahni et al.^7,50^ based on flow cytometry of epitope tagged SLC41A2 heterologously expressed in DT40 chicken cells. Sequence comparisons of lemur, human, and mouse SLC41A2 are shown in Fig. S5. **(c)** Experimental scheme and timeline for constructing human *SLC41A2*^-/-^ iSANCs and analyzing the effects of the gene knockout (see Methods for details). **(d)** Immunofluorescence photomicrographs of wild-type iSANCs co-stained for SLC41A2 (green) and phospholamban (PLN, red), a sarcoplasmic reticulum (SR) transmembrane protein, with DAPI counterstain (blue, nuclei). Insets, close up of boxed region showing a single iSANC. Note colocalization of SLC41A2 with SR-marker PLN. No SLC41A2 staining was detected in SLC41A2^-/-^ iSANCs (Fig. S6c), as expected for the early truncation and frameshift deletion. Scale bar, 10 µm. n=3 technical replicates. **(e)** Functional calcium imaging with Fluo-4 AM of cultured wild-type control and *SLC41A2*^-/-^ iSANCs. Representative traces (left, amplitude normalized to F_peak_/F_0_) showing the rhythmic calcium transients (spikes) of wild-type iSANCs and the slower firing rate of *SLC41A2*^-/-^ iSANCs. Violin plots quantify the effects of *SLC41A2*^-/-^ knockout on calcium transient firing rate, amplitude, time to peak, tau decay time, and duration at 50%. Dark dotted line, median; light dotted lines, quartiles. n= number of cells analyzed (28 WT, 37 *SLC41A2*^-/-^ iSANCs); p-values (two-tailed with Mann-Whitney test): *, <0.05; **, <0.005; ***, <0.0005; ****, <0.00005; ns, not significant. Scale bar for tracings, 1 second, 1 normalized units. **(f)** Functional magnesium imaging with Mag-Fluo-4 AM of cultured wild-type control and *SLC41A2*^-/-^ iSANCs. Representative traces (top, amplitude normalized to F_peak_/F_0_) showing rhythmic magnesium transients of wild-type iSANCs (top left), and three examples of altered waveforms of *SLC41A2*^-/-^ iSANCs (bottom left, ectopic, stunted, and irregular mini-transients; top right, ectopic, stunted transients appearing to fuse with normal transients to produce an irregular rhythm; bottom right, transients with delayed time to peak, tau decay time, and duration at 50%). Irregular rhythm plot quantifies % of iSANCs that manifest mini-transients. Violin plots as above quantify the effects of *SLC41A*2^-/-^ knockout on magnesium transient firing rate, amplitude, time to peak, tau decay time, and duration at 50%. n=number of iSANCs analyzed (27 WT, 30 *SLC41A2*^-/-^ iSANCs) (statistics and significance as above, except p-value for irregular rhythm plot determined by two-tailed Fisher’s exact test). Scale bar for tracings, 1 second, 0.2 normalized units. **(g)** Model of SLC41A2 and magnesium dynamics in human iSANCs, the cardiac pacemaker cells (modified from Weisbrod D et al^54^). Current models for cyclical firing of SAN cells (“automaticity”) invokes two cyclically-active and coupled cellular “clocks”: a voltage membrane clock and an intracellular, sarcoplasmic reticulum-based calcium clock^54–56^. The cardiac cycle begins by hyperpolarization activation of plasma membrane channel HCN4 that initiates the membrane clock (which generates the I_f_ funny current and causes Na^+^ influx, alone or in combination with other Na^+^ influx channels including NACLN, TRPM7, TRPM4, and T-type calcium channels (CaT: CACNA1G and CACNA1H that result in Ca^2+^ influx) and calcium clock (the SR-transmembrane ryanodine receptor, RYR1, which pumps Ca^2+^ from the SR into the cytoplasm and activates the plasma membrane Na^+^-Ca^2+^ exchanger (NCX, encoded by *SLC8*) that in turn exchanges the cytoplasmic Ca^2+^ for Na^+^ into the cell). The resulting increase in membrane potential produced by the coupled clocks activates plasma membrane L-type calcium channels (LTCC), causing a rapid but transient Ca^2+^ influx (calcium transient, gold sunburst) and the upstroke of the action potential (phase 0 depolarization), triggering each cycle of cardiac contraction. Much like the dynamic regulation of calcium, we observed cyclical magnesium transients (blue sunburst) in wild-type iSANCs and slower and broader calcium transients and altered magnesium transient dynamics and waveforms in *SLC41A2*^-/-^ iSANCs. We hypothesize a third clock, the “magnesium clock”, which interacts with and influences the membrane and calcium clocks, and involves the SLC41A2 magnesium transporter and dynamic regulation of the intracellular distribution of Mg^2+^ between the sarcoplasmic reticulum and cytosol. Transient high levels of cytosolic Mg^2+^ counteract the Ca^2+^ and thereby slow the pacemaker firing rate, perhaps through the known ability of Mg^2+^ to directly compete with Ca^2+^ for binding to the key L-type calcium channel (LTCC)^75–78^.

We used functional calcium imaging of cultured iSANCs to examine the effect of the *SLC41A2* knockout on the transient calcium spikes that underlie cardiac pacemaker function. Control wild-type iSANCs fired at ∼100 spikes per minute, whereas the firing rate of the *SLC41A2* knockout iSANCs was ∼20% slower (80 spikes per min) (Fig. 6e). This reduction in firing rate was associated with increased time to peak, increased tau decay time, and increased duration of the peak of the calcium transients, whereas spike amplitude was unaffected. We conclude that *SLC41A2* is required cell autonomously in human iSANCs to maintain their normal rate and waveform of pacemaker firing.

SLC41A2 is one of three members of a small family of multipass transmembrane proteins that function as magnesium-selective transporters^50,51^ (Fig. 6b). The best studied member is SLC41A1, which transports magnesium ions across the plasma membrane. SLC41A3 is a mitochondrial magnesium transporter. SLC41A2 is the least well understood though it too can function as a magnesium ion-selective transporter when expressed in heterologous cells (Xenopus oocytes, chicken DT40 cells)^6,7^. In wild-type iSANCs, SLC41A2 was not detected on the plasma membrane but instead localized to an intracellular compartment, the sarcoplasmic reticulum (Fig. 6d). Magnesium is well established as a gatekeeper for calcium entry and cardiac contraction^52^, but its role in pacemaker cells and cardiac pacemaking has not been previously investigated. To explore the impact of SLC41A2 on magnesium levels and dynamics in cardiac pacemaker cells, magnesium functional imaging was performed on human iSANCs, as done above for calcium. We discovered that magnesium levels are not constant in wild-type iSANCs but rather they spike regularly, much like the canonical calcium spikes (Fig. 6f). However, in *SLC41A2* knockout iSANCs, the magnesium transients occurred at irregular intervals, sometimes with small ectopic spikes, and with overall increases in magnesium amplitude, time to peak, tau decay time, and peak duration. Thus, magnesium spikes regularly in human iSANCs, and *SLC41A2* is required for the proper timing, waveform, and amplitude of these intracellular magnesium transients.

## DISCUSSION

We established a primate model for cardiac arrhythmias through systematic ECG screening of over 350 mouse lemurs that identified eight naturally-occurring arrhythmias resembling human pathological conditions plus seven other variants seen in potentially pathological human conditions. The most common arrhythmia (2.2% prevalence) was a profound, intermittent bradycardia resembling human SSS. Lemur SSS is an autosomal recessive and fully penetrant disorder that can be detected early in life and have significant clinical sequelae associated with cardiac chronotropic incompetence including sudden death, like the human disease if left untreated. Genome sequencing and linkage analysis of the lemur pedigree mapped the disease locus to a 1.4 Mb interval on chromosome 7 containing nine candidate genes with linked polymorphisms. The most appealing candidate is *SLC41A2*, a little studied transmembrane magnesium transporter whose mouse knockout phenotype is cardiac-selective like the lemur disease but manifests as a ventricular conduction defect^8^ (wide QRS complex) rather than the sinoatrial node (SAN) pacemaker dysfunction of lemur and human SSS. SLC41A2 is expressed in iPSC-derived human SAN cells (iSANC) and localizes to the sarcoplasmic reticulum, and CRISPR-Cas9 mediated *SLC41A2* knockout slowed the rhythmic calcium transients that underlie their automaticity and pacemaker function. Thus, *SLC41A2* is a critical new component of the human pacemaker and functions cell autonomously to increase calcium spike frequency and pacemaker activity. Other SAN ion transporters and channels such as HCN4 are also critical for calcium spikes and pacemaker activity^53,54^, and when they are mutant cause human SSS^46,47^; this supports our provisional assignment of *SLC41A2* as the lemur SSS disease gene and suggests it functions similarly in the lemur but not the mouse cardiac pacemaker. Likewise, our results nominate human *SLC41A2* as a candidate gene for familial forms of human SSS not attributable to a canonical SSS gene such as *HCN4*.

Our results also revealed rhythmic magnesium transients in human iSANCs, like the classical calcium spikes. The magnesium spike pattern, level and waveform were altered in *SLC41A2*-knockout iSANCs, presumably due to altered magnesium flux from the sarcoplasmic reticulum normally mediated by the SLC41A2 transporter. These results lead us to propose that SLC41A2 and intracellular magnesium dynamics comprise an important but previously unappreciated axis of the cardiac pacemaker. Current models of pacemaker automaticity invoke a coupled-clock system operating within each SAN cell^54–56^ (Fig. 6g). A plasma membrane oscillator or clock (“membrane clock”), mediated by a set of time and voltage-dependent membrane ion channels including HCN4, is coupled to a sarcoplasmic reticulum calcium clock (“Ca clock”). The latter stimulates sodium influx in exchange for calcium gated by the Ryanodine Receptor (RyR), working in tandem with the membrane clock to depolarize the nodal cell membrane and activate the voltage-gated L-type calcium channel (LTCC), resulting in the rhythmic calcium spikes and firing of each heartbeat-inducing action potential. Our results identify a third oscillator (“magnesium clock”) in each iSANC that appears to interact with and regulate at least one of the other two clocks. Although the mechanisms by which magnesium transients are generated and impact the other clocks remain to be defined, this new oscillator must involve additional channels and transporters beyond SLC41A2 because magnesium transients persisted in *SLC41A2* knockout iSANCs, albeit with altered kinetics and waveforms. Magnesium has long been known to influence cardiac automaticity and conduction^52^, and magnesium therapy is used clinically to treat or stabilize certain human arrhythmias^57–59^. The identification of SLC41A2 function and magnesium oscillations in iSANCs provide a rationale and target for development of new, mechanism-based therapeutics for SSS and other sinus node arrhythmias. The current standard of care for SSS is placement of an electronic pacemaker, and SSS is the most common indication for pacemaker placement^44^. A medical therapy would spare patients the invasive procedure and clinical complications of the implanted device.

Our systematic screen and genetic mapping of a heritable trait establish mouse lemurs as a tractable genetic model organism. Their exceptionally small size (60 gm), short generation time (6-8 months), large litter size (typically 2-3), and modest lifespan (∼6 years) for a primate, and the relative ease in creating, maintaining, and expanding a laboratory colony^5^, facilitated our systematic phenotypic screening, multigeneration pedigree analysis, and genetic mapping -- classical genetic approaches impractical for other non-human primates. By focusing our screen on easy to identify and medically important traits (arrhythmias) for which there is urgent need for new preclinical models, we were able to uncover natural models for many significant human arrhythmias. Most importantly, we demonstrated the value of this new genetic model by discovering a new gene (*SLC41A2*), molecules (SLC41A2, magnesium ions), and mechanism (magnesium transients) of the primate pacemaker, and by identifying a novel candidate gene and therapeutic target for human sinus node arrhythmias.

These forward genetic approaches can now be applied to any familial trait in mouse lemur, including atrial fibrillation (Fig. S2) and other cardiac, oncologic, and motor pathologies identified in our deep phenotyping and transcriptomic profiling screens. These complement the reverse genetic approach that we also recently established, in which natural knockout mutations in lemur genes, including three of the hundreds of primate genes missing in mice, have been identified by genome sequencing and their transcriptional effects described^60^, and the cellular and molecular biology approaches enabled by an extensive transcriptomic cell atlas of over 750 cell types across 27 organs^49,60^. We hope these genetic, cellular, and molecular advances for mouse lemur provide the experimental foundation to achieve new insights and ultimately interventions for many primate diseases and other aspects of primate biology, behavior, and ecology.

## ACKNOWLEDGMENTS

We are grateful to all of our hosts, guides, and mentors at the CNRS-MNHN (Museum National d’Histoire Naturelle) in Brunoy, France and the Centre ValBio (CVB) field station at the Ranomafana National Park in Madagascar for their contributions to the longitudinal phenotyping of mouse lemurs. We thank Zicheng Zhao for help with genomic DNA purification; Vida Shokooshi and John Coller (Stanford Genomics Center) for input on DNA library preparation and high-throughput sequencing; Ramesh Nair (Stanford Genetics Bioinformatics Service Center) for help with establishing a workflow for DNA variant calling; the Stanford Research Computing Center for computational resources and support including Stanford Sherlock and Oak clusters; and Maria Peterson for figure preparation and administrative support. We thank Richard Lewis for valuable discussions on calcium and magnesium dynamics; Ross Metzger, Sori Jang, Christin Kuo, and members of the Krasnow lab for helpful discussions and comments on the manuscript; and Eldrin Lewis and the Stanford Division of Cardiovascular Medicine leadership for their encouragement and support. S.C. was supported by the Advanced Residency Training at Stanford (ARTS) program, C.J.K. by a France-Stanford Center for Interdisciplinary Studies Grant and a Stanford Child Health Research Institute Postdoctoral Award, and L.R. by an American Heart Association Postdoctoral Award. This work was supported by funding from the Howard Hughes Medical Institute and the Vera Moulton Wall Center for Pulmonary Vascular Disease (M.A.K.). M.A.K. is an investigator of the Howard Hughes Medical Institute.

## METHODS

### Animals

Laboratory *M. murinus* lemurs were from the closed captive breeding colony at the Muséum National d’Histoire Naturelle in Brunoy, France. Lemur husbandry and care has been described^38^. Briefly, lemurs were housed indoors individually or in small groups in cages enriched with tree branches, wooden nest boxes, and artificial lighting. Photoperiod light cycles were alternated from 14:10-h (light) to 10:14-h (dark) every six months to stimulate photoperiod-dependent seasonal behavior and metabolic changes, and ambient temperature was maintained at 24-26°C and humidity at ∼55%. Diet included cereal, fresh fruit, gingerbread, milk, and eggs, and water was provided *ad libitum*. Regular health monitoring included weekly assessments and monthly veterinary examinations. All procedures adhered to European ethical regulations for animal use in biomedical research and were approved by the Animal Welfare board of UMR 7179. Seven descendants of this breeding colony, used only for whole genome sequencing analysis, were transferred to and maintained at the Stanford Animal Research Facility at Stanford Medical School for noninvasive phenotyping and genetic research as approved by the Stanford University Administrative Panel on Laboratory Animal Care (APLAC #27439) and in accordance with the Guide for the Care and Use of Laboratory Animals, as detailed previously^49,61^.

Wild *M. rufus* and *M. nova* lemurs living in and near Ranomafana National Park (RNP) that have been followed longitudinally in the field for decades by serial capture-and-release^5,62,63^ were analyzed at Centre ValBio (CVB) field station, which houses a modern genetics and molecular biology laboratory at the edge of RNP^64^. Briefly, individual lemurs were captured in live rodent traps baited with a banana slice and secured to a tree branch. Traps with lemurs were brought to the lab for lemur deep phenotyping including ECG analysis, and the lemur was then released back into the wild at the capture site several hours later. Lemurs were named and identification microchips placed subcutaneously the first time they were captured and phenotyped.

### Electrocardiography and heart rate analysis

Single-lead, 3-lead, and derived 6-lead point-of-care electrocardiography (ECG) assays were based on the Kardia system (AliveCor®), an FDA-cleared device for human ECG detection that we adapted for application to mouse lemur. For single-lead recordings, the KardiaMobile® device with two electrode pads (each designed to contact one or two fingers of a human hand) was remotely connected to a smartphone (Apple iPhone models 8, XS Max, 12 Pro, and 14) via the Kardia software app (versions 5.27.0 and 5.41.0). For 3-lead and derived 6-lead recordings, the KardiaMobile® 6L device, which has a third electrode pad on the back of the device that is designed to contact the skin of the human left leg, was adapted by removing the back plate and soldering a wire to the conductive post of the back pad. The device was then mounted in a square (9.3 cm tall, 10 cm wide, 1.2 cm thick) 3D-printed polylactic acid holder with a copper plate (2.9 cm x 3.2 cm) attached on top with a conductive adhesive (Fig. 1c), and the wire from the back post was then soldered to the copper plate to complete the circuit. This provided three electrode pads on top of the modified device, the two standard pads (one for each hand of the lemur) and the added copper pad (for the left foot).

Kardia software and hardware were optimized to reliably and accurately capture the mouse lemur heart rate and cardiac cycle, initially using the single- and 3-lead systems, and then validated with the derived 6-lead system. Unlike rodents, for which ECG recording typically requires anesthesia, the lemurs were docile and required only gentle support on the ECG platform, such that their arms rested on the device with one hand on each pad, and their back left leg guided onto the copper pad. ECGs were recorded for 10-30 seconds. ECGs were repeated sequentially as needed (up to 3 times) to facilitate desensitization of the subjects to the test. Serial recordings were generally consistent, but if the first traces were different they were discarded. Any differences observed were primarily from artifacts introduced by dirty hands of the subject that caused poor electrode contact, reducing and distorting the ECG signal. Recordings were transmitted remotely to the Kardia smartphone App, with ECG raw data available online for cardiologist review and analysis. Automated ECG analysis software designed for human ECGs (Kardia App) did not reliably interpret lemur ECG traces, so ECG reading and pathology calls were done manually by two or three board-certified cardiologists (S.C., V.F., D.L.) for each trace. Some individuals were studied longitudinally. ECGs were performed from 2016-2024.

Heart rate ranges for each of the three Microcebus species were determined by calculating 95% confidence intervals (CI) using 2 standard deviations from the mean. Values falling below the lowest limit of the 95% CI defined the range for sinus bradycardia. For sinus tachycardia (ST), the 95% CI was calculated using only ECGs interpreted as ST (T-wave visibly fused with preceding QRS-complex). Values below the lowest limit of this 95% CI for ST (but above the highest cutoff for sinus bradycardia) was taken as the heart rate range for normal sinus rhythm.

### Lemur family pedigrees

The pedigrees of the laboratory *M. murinus* lemurs with sick sinus syndrome (SSS) (Fig. 4) and atrial fibrillation (Afib)/premature atrial contractions (PACs) (Fig. S2) were constructed from breeding records of the colony and DNA genotyping of selected, highly polymorphic loci. The pedigree of the full colony is under construction, and experimental protocol and full set of results will be detailed elsewhere (C.K., J.P., C.R., A.A., J.T., et al, unpublished data). Briefly, maternal lineage and siblings were obtained directly from the breeding record, which includes the identities of the mother and each offspring (typically 2-3) in the litter. However, in most cases paternity was ambiguous in the record because for mating a female lemur was usually bred with up to 4 adult males, any one of which could have fathered the litter. Maternity was confirmed and paternal ambiguity was resolved by isolation of genomic DNA from each individual, its mother, and all potential fathers, and polymerase chain reaction amplification and DNA sequencing of five selected, highly polymorphic loci (Mm03, Mm09, Mm10, Mm21, Mm39)^65,66^. In almost all cases, comparison of the sequences was sufficient to identify the biological father and confirm the identity of the mother, as shown in the SSS and Afib/PAC pedigree figures (Fig. 4, Fig. S2). In the rare cases in which genomic DNA was not available for genotyping, or the genotyped loci did not fully resolve paternity, paternity is reported as “?#” in the figure.

### Whole-genome sequencing

Whole blood (150-250 ul) was obtained from 35 *M. murinus* lab animals by saphenous vein pricks with a small gauge (22 or higher) needle and collected by capillary action into microfuge tubes. Genomic DNA (gDNA) was extracted from each blood sample using Qiagen DNeasy Blood & Tissue Kit (#69504) as described by the manufacturer, and gDNA concentration was determined by fluorimetry (Invitrogen Qubit^TM^ 3 Fluorometer, 1X dsDNA high sensitivity assay kit Q33230). Quality control to ensure absence of extensive DNA fragmentation was done by gDNA electrophoresis on 3% agarose gels. gDNA (1 ng) was used for sequencing library preparation with dual unique i5 and i7 indexing primers for the Illumina platform as described^49^; no significant differences in tagmentation size, library quality, and sequencing read depth were observed between libraries prepared with our in-house protocol versus the commercial Illumina kit. gDNA libraries were sequenced to achieve saturation on a NovaSeq 6000 System (Illumina) using 2 x 150 bp paired-end reads and 2 x 10 bp dual unique index reads with 300 cycle kits designed for either the S2 or S4 flow cell configuration (Illumina, 20091658, 20028314, 20028312). Sequencing data was outputted in FASTQ format and sequencing quality assessed with FastQC (v0.11.9).

### Sequence variant identification and segregation in SSS pedigree

The FASTQ DNA sequencing files for the 35 *Microcebus murinus* lemurs were aligned to Mmur 3.0 genome assembly (NCBI RefSeq assembly accession: GCF_000165445.2; indexed using Burrow-Wheeler Aligner (version BWA-0.7.17 r1188); NCBI *Microcebus murinus* Annotation Release 101, GTF format) and sequence variants identified using Sentieon® (software version 202112.01) DNAseq® and germline variant calling pipeline^67^. Mapping, alignment quality metric determination (QualiMap(v2.2.1), removal of base duplicates, base quality score recalibration (BQSR), and variant identification were carried out individually for each lemur using the customized options and computational scripts available on Github (https://github.com/scthree/SSS). Individual variant calling files were then joint genotyped together by a coalescent model to produce a single, multi-sample variant calling file. Methods for identification and mapping of SNP/indel variants that co-segregated with SSS in the lemur pedigree are detailed below; the full *M. murinus* SNP/indel library created from these 35 individual genomes will be described elsewhere.

A database of the *M. murinus genome* (Mmur3.0 assembly and annotation files) was constructed for the SnpEff & SnpSift toolbox (v4.5)^68^, and then the toolbox used (java version 11.0.11, detailed on Github, https://github.com/scthree/SSS) to annotate and predict the functional effect of the identified SNP/indel variants in the joint genotyped variant calling file described above. SnpSift was then used to filter for variants that co-segregated throughout the pedigree with the assigned phenotype of each individual (SSS-affected, carrier, wild-type) in a simple Mendelian autosomal dominant inheritance pattern, or an autosomal recessive fashion, assuming full penetrance of the phenotype (https://github.com/scthree/SSS). The SSS segregation pattern in the pedigree excluded X-linked and mitochondrial inheritance patterns, so these inheritance patterns were not further investigated here.

### Creating human cardiac sinoatrial nodal pacemaker cells (iSANCs) from human iPS cells (iPSCs)

Differentiation of human iSANCs from iPSCs was done as described^69^ with the following modifications. Wild-type human female iPSCs (clone #273, Stanford Cardiovascular Institute iPSC Biobank; https://med.stanford.edu/scvibiobank/request-cells.html) were cultured in 5% CO_2_ incubators at 37 °C in three stages (A, B, and C, 5 days each over 15 days, instead of 3 days each over 9 days as previously described) to promote SANC differentiation (Fig. 6c). Starvation of cells was carried out through SANC differentiation and purification (∼day 50, instead of only through day 30 (stage B) as previously described) to enrich for nodal cells (fatty-acid metabolism) over iPSCs (anaerobic metabolism); hence, glucose and insulin were not added to the base culture medium until ∼day 51 (beginning of “SANC maturation”), which was then used for the remainder of the protocol: Roswell Park Memorial Institute, RPMI 1640 medium with glucose (Gibco, Thermo Scientific #11875093) supplemented with 10% B-27 containing insulin (Gibco, Thermo Scientific #12587010), glutamine substitute GlutaMAX (at 2 mM, Gibco, Thermo Scientific #35050061), 1x Minimum Essential Medium, MEM non-essential amino acids (Gibco, Thermo Scientific #11140050), and 1x penicillin-streptomycin (Gibco, Thermo Scientific 15070063). During SANC purification and maturation, cells were washed (in phosphate buffered saline, PBS, pH 7.4) and culture media replaced every 2-3 days. Mature SANCs were identified by their morphology (small, spindle-like cells forming clusters for functional synchronization) and beating rate (∼100 beats per minute) under light microscopy.

### Generating *SLC41A2^-/-^* knockout human iSANCs with CRISPR/Cas9

CRISPR/Cas9 single guide RNAs (sgRNAs) targeting the N-terminal coding region in an early common exon (exon 2) of human *SLC41A2* were bioinformatically designed using CRISPR design tools from Synthego^70^, Integrated DNA Technologies (IDT)^71^, and CRISPick (Broad Institute)^72^. Three customized sgRNAs (Fig. S6a) were selected for their predicted high targeting efficiency and specificity (low off-target effects). The *SLC41A2* targeting sgRNAs (3 nmol each) were synthesized (Synthego), and a pooled sgRNA (‘multi-guide’) working stock solution (30 µM) was prepared and used along with similar working stock solutions of sgRNA controls (positive control, multi-guide sgRNAs targeting human TRAC, 14637241-2; negative control, scrambled synthetic sgRNA #1, 4637241-3) according to the manufacturer’s instructions (Gene Knockout Kit v2, 14637241-1; transfection optimization kit 14637241-2). CRISPR/Cas9 editing was carried out by a proprietary protocol (Neon Electroporation, Synthego). Briefly, on day 1 of the protocol, human female iPSC clone #273 cells were cultured in a humidified chamber under 5% CO_2_ at 37℃ in a serum-free complete iPSC cell culture medium (mTeSR media, Stemcell Technologies) in Matrigel-coated plates (Corning 354603). Ribonucleoprotein complexes were assembled by combining sgRNA:Cas9 in a 9:1 ratio (from 30 µM multi-guide sgRNA stock solution and 20 µM SpCas9 2NLS nuclease stock solution, to make final working solutions of 12.9 µM and 0.07 µM, respectively in proprietary resuspension buffer R). On culture day 3, when iPSCs were ∼70% confluent, cells were dissociated, counted, and 1-2 x 10^6^ cells were centrifuged (500 g, 5 min, 4℃) in a sterile microfuge tube, washed with PBS, and resuspended in 50 µL of resuspension buffer R. For each transfection, RNP solution (7 µL) was added to 5 µL of cell suspension, and transferred into a chilled cuvette for electroporation (Bio-Rad Gene Pulser Xcell system, two pulses, 1.4 kV, 40 ms pulse width). Electroporated cells were immediately transferred to one well of a pre-warmed (37℃) Matrigel-coated 96-well plate (Corning 3603) containing 100 µl of mTeSR media and recovered for three days at 37℃ in a humidified 5% CO_2_ incubator. On day 7, an aliquot of the recovered cells was harvested, and genomic DNA was isolated, the editing site amplified by PCR, and the PCR product sequenced (Sanger sequencing) to assess editing efficiency. Limit dilutions were made from the remaining cells and seeded at the following densities in mTeSR, 0.25 cells / 100 µL, 0.5 cells / 100 µL, 1 cell / 100 µL, in three 96-well plates, respectively. Single colonies were then picked from wells in which only one colony had grown (day ∼14), ensuring isolated cells had originated from a single cell; wells with two or more colonies were discarded. When clonal colonies reached ∼100-300 µM diameter (day ∼17), they were passaged into 24-well plates (Corning 3524). After colonies reached ∼70% confluence (∼day 20), *SLC41A2^-/-^* knockout was confirmed by PCR analysis on agarose gel (∼320 bp amplicon vs. ∼440 bp in wild-type), Sanger sequencing (Fig. S6b), and immunofluorescence staining (Fig. S6c), and these were selected for subsequent differentiation into iSANCs as described above. iSANCs differentiated from non-CRISPR-electroporated iPSCs were used as wild-type *SLC41A2^+/+^*controls in the immunofluorescence and functional calcium and magnesium imaging experiments. *SLC41A2^-/-^* mutant and wild-type control iSANCs are available on request.

### Immunofluorescence analysis of human iSANCs

Primary antibodies and dilutions used were SLC41A2 rabbit polyclonal antibody (Invitrogen, Thermo Scientific PA558483; 1:50 dilution), PLN mouse monoclonal antibody (Invitrogen, Thermo Scientific MA3-922; 1:50), and fluorescent secondary antibodies were donkey anti-rabbit Alexa Fluor Plus 488 (Invitrogen, Thermo Scientific A32790; 1:200) and donkey anti-mouse Alexa Fluor Plus 555 (Invitrogen, Thermo Scientific A32773; 1:200). For immunostaining, human *SLC41A2^+/+^* and *SLC41A2^-/-^*iSANCs were plated in 35 mm glass bottom dishes and fixed in 4% PFA (pH 7.4) at room temperature (RT) for 20 min. After washing three times with sterile PBS, cells were permeabilized in 0.1% Triton X-100 in PBS and incubated at RT for 25 min. After another round of washing, samples were blocked in 2% BSA in PBST and incubated at RT for 2 hours, washed again, and subsequently incubated in 2% BSA/PBST with primary antibody at 4℃ overnight. After washing, cells were incubated with fluorescent secondary antibody and DAPI (Thermo Scientific 62248; 1:500) in 2% BSA/PBST for 2 hours at RT protected from sunlight. After a final wash, cells were mounted in an antifade mounting media (Vectashield® Plus H-2000) and imaged by confocal fluorescence microscopy (Leica Stellaris 8).

### Functional calcium and magnesium imaging of human iSANCs

Live calcium imaging of human iSANCs was performed using cell permeant calcium indicator Fluo-4 AM (Thermo Scientific F14201) as described^73^. Briefly, iSANCs were seeded for imaging at a density of 500,000 cells/mL in 10% KOSR RMPI/B27 media (KnockOut Serum Replacement, Thermo Scientific 10828-028; RMPI 1640 Medium, Life Technologies 11875-119; B27 supplements, Life Technologies 17504-044). Cells were loaded with 5 µM Fluo-4 with 0.1% Pluronic F-127 (Thermo Scientific P3000MP) for 10 min at RT in Tyrode’s solution (140 mM NaCl, 1 mM MgCl_2_, 5.4 mM KCl, 1.8 mM CaCl_2_, 10 mM glucose, and 10 mM HEPES, pH 7.4 with NaOH). Ca^2+^ handling was observed using an inverted microscope (Zeiss LSM980) mounted with a 63x oil immersion objective. During experiments, cells were maintained at 37 °C and constantly gassed with 95% O_2_/5% CO_2_. Line-scan images across the whole cell were obtained to quantify local Ca^2+^ signals with 488 nm excitation and 510 nm emission from the iSANCs. The same protocol was used for magnesium indicator Mag-Fluo-4 AM (Thermo Scientific M14206), with fluorescence excitation at 493 nm and emission collection at 517 nm.

## Data and code availability

All coding scripts and computational options used are available on Github (https://github.com/scthree/SSS).

## SUPPLEMENTARY MATERIALS

**Figure S1.**
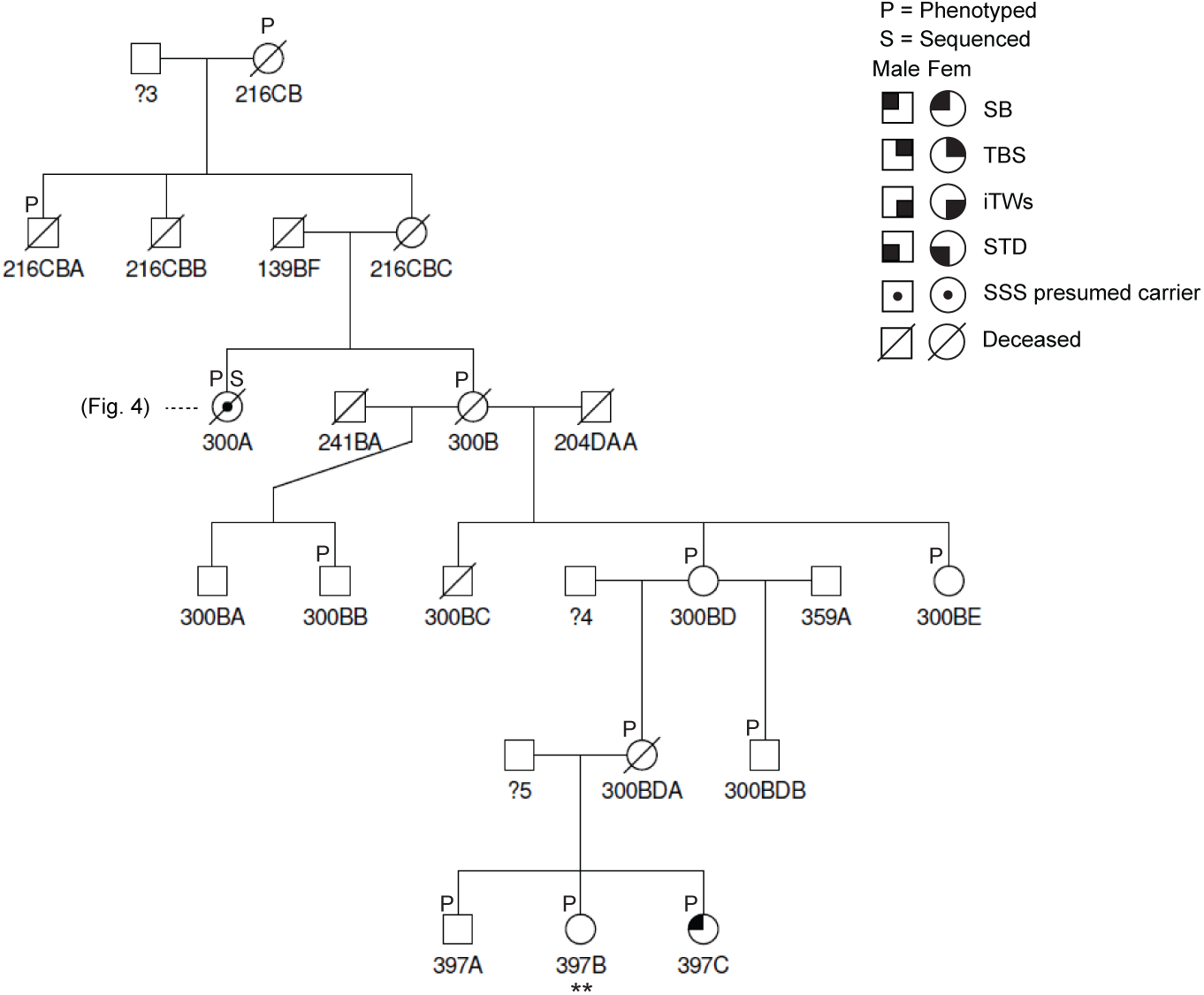
Extended lemur sick sinus syndrome pedigree. Extension of the lemur sick sinus syndrome (SSS) pedigree (Fig. 4) showing the relationship to the seventh affected mouse lemur (397C, diagnosed at 1.5 yrs). Note the relationship between 397C and 300A, her great great aunt and an SSS-carrier in the main SSS pedigree (Fig. 4). Further pedigree construction and genome sequencing are required to resolve the relationship between the primary SSS pedigree and 397C in this extended pedigree. ?3, ?4, ?5, paternities not yet resolved (see Methods). His sister (397B, **), was found to have decreased grip strength at age 1.5 yrs.

**Figure S2.**
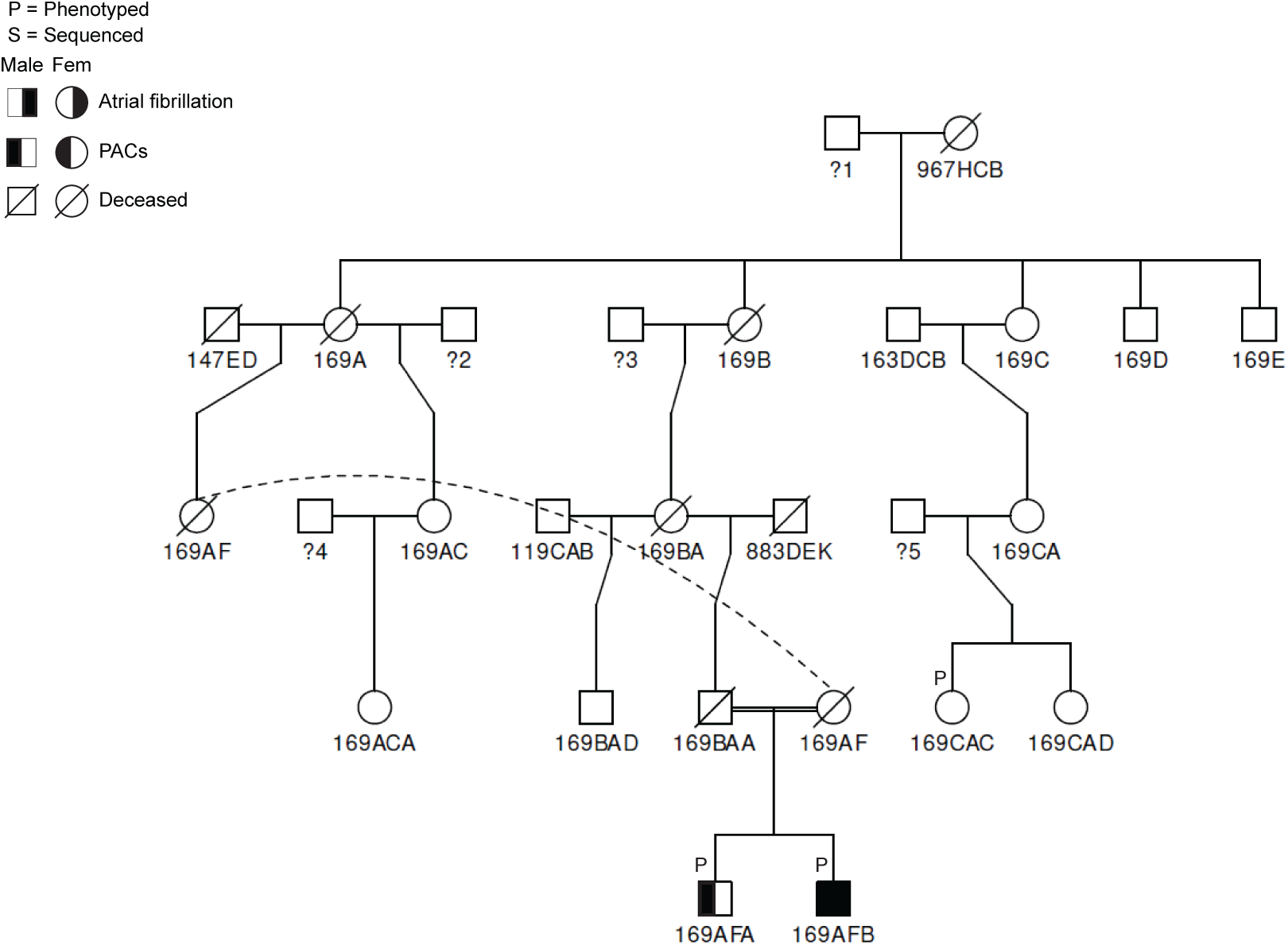
Lemur family with premature atrial contractions and atrial fibrillation. Two of the lemurs diagnosed with premature atrial contractions (PACs) are brothers: 169AFA (diagnosed at age 2.6 yrs) and 169AFB (at age 3.1 yrs). 169AFB was subsequently diagnosed with paroxysmal atrial fibrillation (Afib, at age 4.6 yrs). In humans, PACs are a known precursor to Afib.

**Figure S3.**
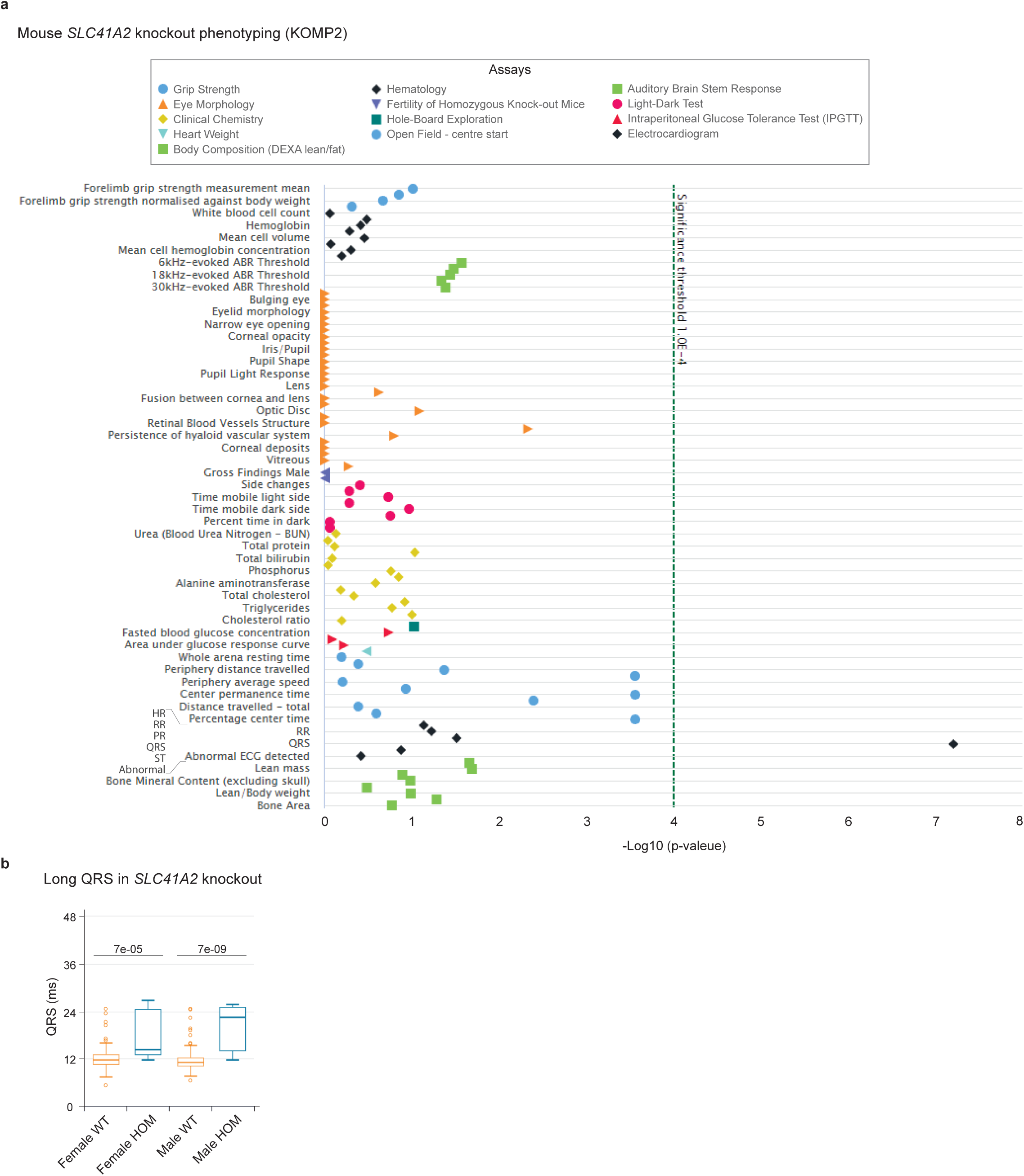
Mouse *SLC41A2* knockout deep phenotyping by KOMP2. (Top) Results of deep organ-wide phenotyping of *SLC41A2*^-/-^ mice (C57BL/6NJ-*Slc41a2^em1(IMPC)J^*/Mmjax) generated and analyzed by the Knockout Mouse Phenotyping Program (KOMP2) at The Jackson Laboratory^8,79^ and reproduced here. The *SLC41A2*^-/-^ knockout is a CRISPR/Cas9-generated 467 bp deletion encompassing *SLC41A2* exon 5 and a flanking 322 bp intronic region that includes the splice donor and acceptor. Deep phenotyping of 348 *SLC41A2*^-/-^ mice by the assays indicated surveying >50 traits revealed a single abnormality: an adult cardiovascular conduction system defect manifesting as a widened QRS complex on ECG (lone black diamond on right). Subsequent phenotyping has also detected isolated skin and vocalization abnormalities^8^. Note that no other ECG abnormalities (black diamonds: heart rate, HR; RR interval, RR; PR interval, PR; ST segment, ST; abnormal rhythms, Abnormal ECG detected) were detected; the absence of HR and RR interval abnormalities are notable here because they indicate normal SA node pacemaker function in the *SLC41A2* knockout. (Bottom) Quantification of the QRS interval in ECGs of adult *SLC41A2* wild-type (WT) and homozygous *SLC41A2*^-/-^ knockout (HOM) female and male mice. n=164 female WT, 169 male WT, 10 female HOM, 5 male HOM; p-values (two-tailed Fisher’s exact test). ms, milliseconds.

**Figure S4.**
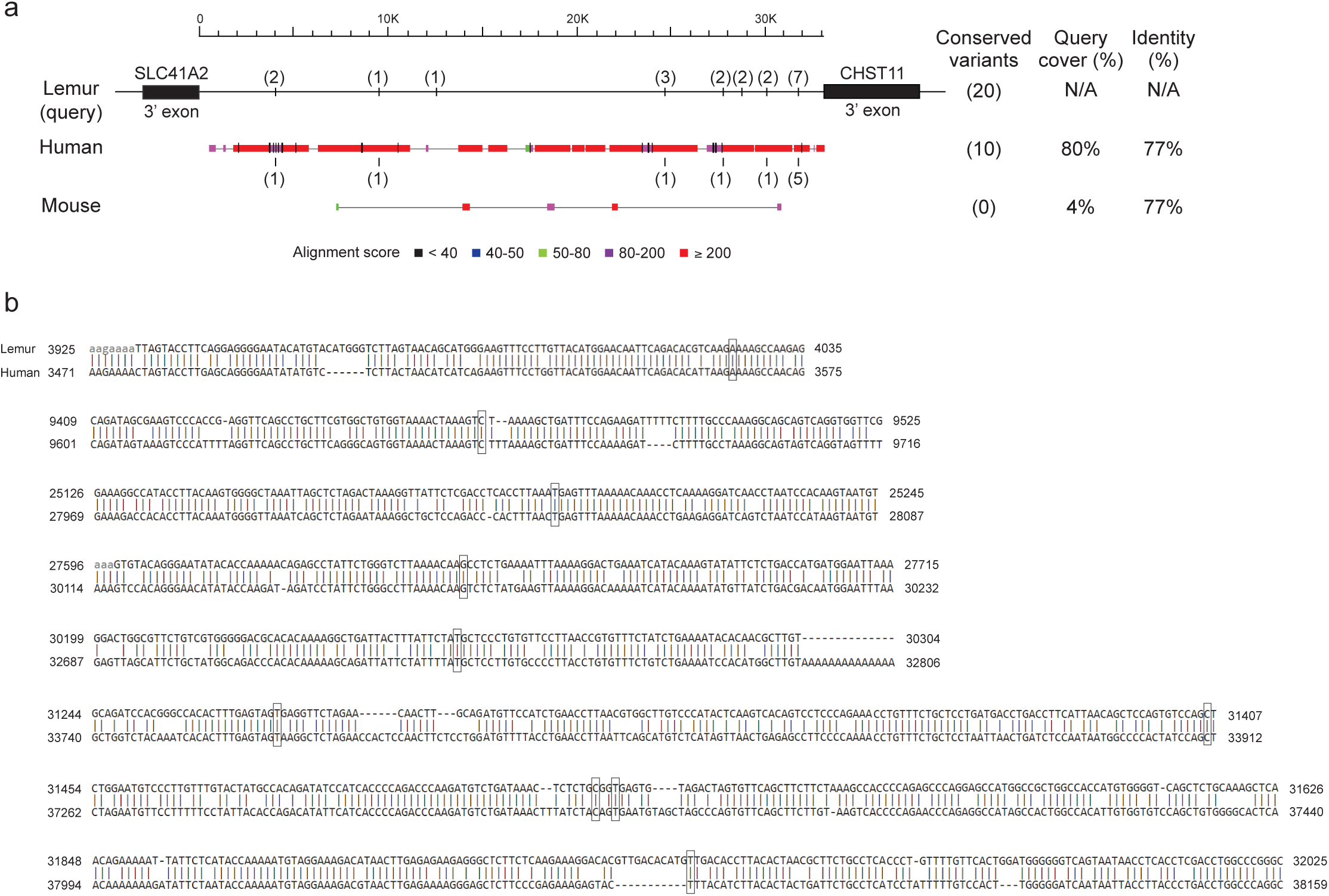
DNA sequence conservation between lemur, human, and mouse in the *SLC41A2-CHST1* intergenic region. **(a)** Cross-species DNA sequence comparison using NCBI nucleotide discontiguous Mega BLAST+ (v. 2.16.0) of the lemur *SLC41A2-CHST11* intergenic region (33,108 bp query, top, see Fig. 5d) with the corresponding intergenic region of human (39,785 bp) and mouse (34,972 bp). The positions of the 20 lemur SSS-linked variants are indicated in parentheses above the line representing the lemur intergenic region, and the 10 that are conserved in humans are similarly indicated below the bar representing the human intergenic region. The mouse intergenic region is divergent from the primates, and none of the 20 SSS-linked variants are conserved. Query Cover, proportion of the lemur intergenic sequence that aligned to the human or mouse intergenic sequence. % Identity, percentage of base pairs that are identical within the aligned regions. Colors of the bars, discontiguous Mega BLAST alignment scores for the corresponding regions (higher scores represent better alignment). **(b)** Lemur vs human nucleotide sequence and conservation (vertical lines) surrounding each of the 10 conserved SSS-linked variants (boxed). Numbers (at left), positions of the sequence shown in the respective intergenic region.

**Figure S5.**
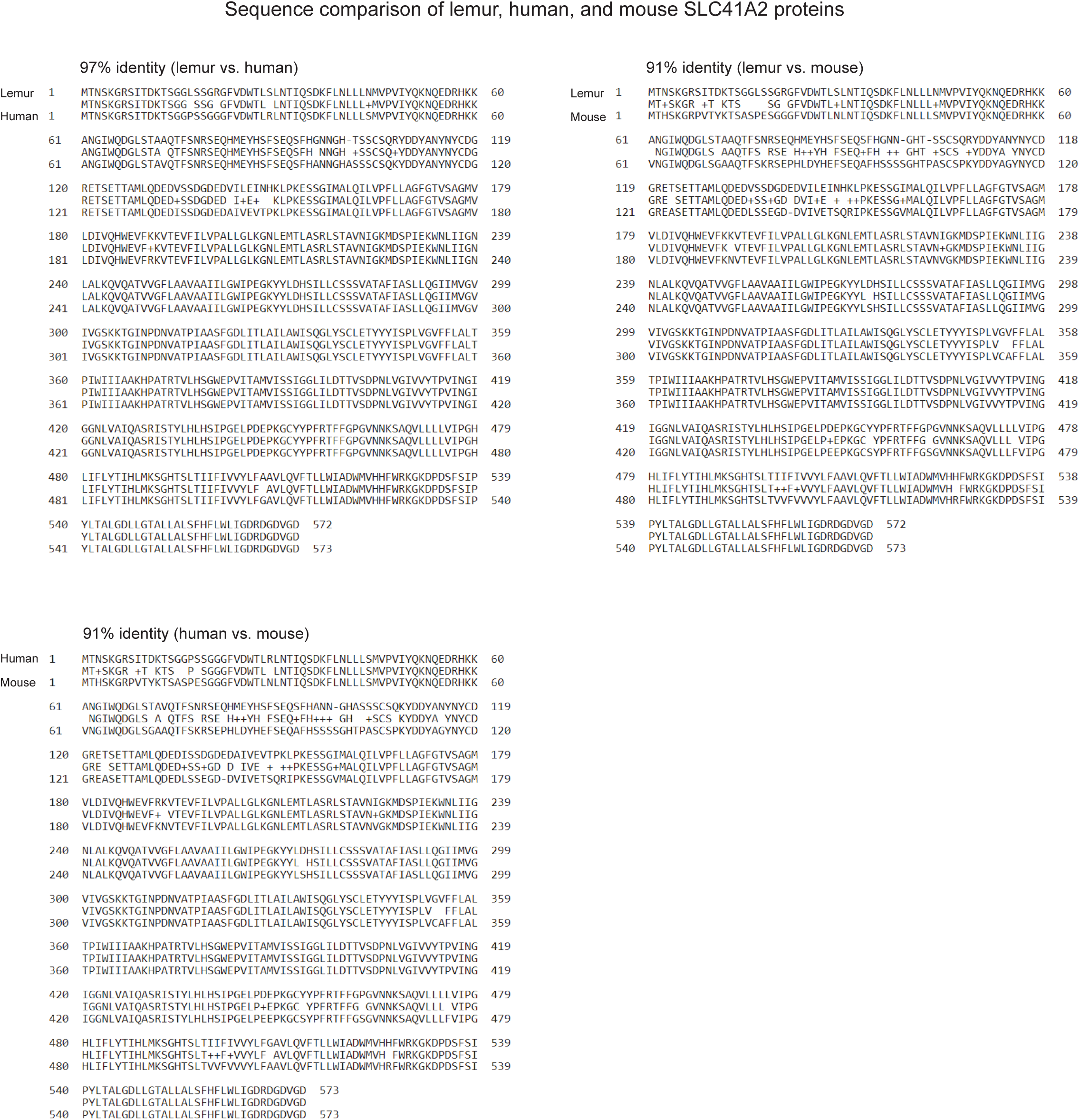
Pairwise cross-species alignments (NCBI protein blastp, BLAST+ v. 2.16.0+) of SLC41A2 protein sequences of lemur, human, and mouse. Values above each alignment show percent amino acid identity.

**Figure S6.**
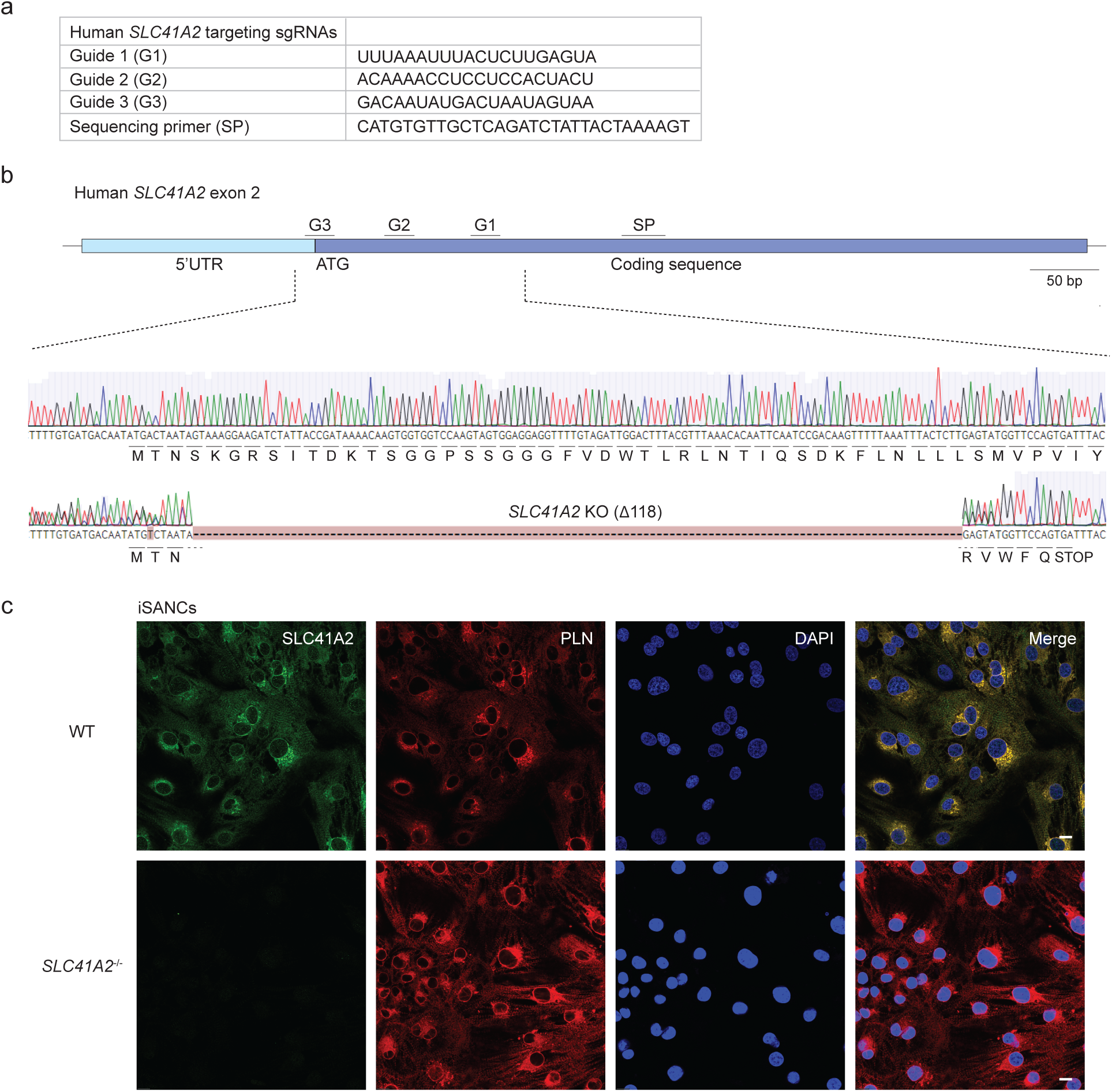
Structure and confirmation of CRISPR/Cas9 knockout of human *SLC41A2*. **(a)** Nucleotide sequences of the sgRNAs (“multi-guide”) designed and used to target human *SLC41A2* for gene knockout in iPSCs (Fig. 6a,c), and the Sanger sequencing primer (SP) used to verify the deletion. **(b)** Map of the first coding exon (exon 2) of human *SLC41A2* showing the hybridization positions of each sgRNA (G1, 2, 3) and the sequencing primer (SP). Light blue, portion of the 5’-untranslated region (5’UTR) in exon 2. Dark blue, coding sequence. Enlarged region below shows Benchling alignment (Multiple Alignment using Fast Fourier Transform, MAFFT v. 7.505) of amplicons generated by Sanger sequencing of genomic DNA harvested from control WT iSANCs (similarly derived from parental wild-type iPSCs) and the *SLC41A2*^-/-^ iSANC knockout clone created and used in this study. Note CRISPR/Cas9 targeting resulted in a 118 bp deletion in exon 2 of *SLC41A2*^-/-^ iSANCs, producing a coding sequence deletion and frameshift at codon 4 resulting in a translation termination codon (STOP) five codons later (residue 9). **(c)** Fluorescence micrographs of control wild-type (top) and *SLC41A2*^-/-^ knockout (bottom) iSANCs co-stained for SLC41A2 (green) and sarcoplasmic reticulum marker PLN (red), with DAPI counterstain (nuclei, blue). SLC41A2 staining is absent in knockout mutant cells. Scale bar, 10 µm. n=3 technical replicates.

**Table S1.** Major differences between mouse and human cardiac conduction systems. Major differences in anatomy, molecular biology, genetics, biochemistry, and physiology between the cardiac conduction systems of humans and mice. ECG, electrocardiogram; SVC, superior vena cava; RV, right ventricle; LV, left ventricle; QRS, QRS complex (representing ventricular depolarization) on ECG.

| Conduction System Feature | Mice | Human |
| --- | --- | --- |
| Sinoatrial node location | Superior vena cava / Right atrium | Right atrium |
| Atrioventricular node | Less distinct | Distinct |
| His-Purkinje system | Less distinct and organized, fewer fibers | Distinct and organized |
| Superior vena cava (trigger for arrhythmias) | Two (retained left SVC) | One |
| Pulmonary veins (site of origin for arrhythmias) | Converge to a solitary opening in left atrium | Four distinct openings in left atrium |
| K<sup>+</sup> channels for repolarization | *KCNA5, KCNB1, KCNKA/KCNB3* | *KCNQ1, KCNH2* |
| Sinoatrial node action potential | No clear plateau phase, rapid repolarization | Defined plateau phase, slower repolarization |
| T-wave on ECG | Not well resolved, merged with QRS | Discrete from QRS |
| Ventricular depolarization pattern | RV -> LV or LV apex -> RV | Interventricular septum -> both ventricles |
| Heart rate | 600-800 bpm | 60-100 bpm |

Abbreviations: ECG, electrocardiogram; SVC, superior vena cava; RV, right ventricle; LV, left ventricle; QRS, QRS interval on electrocardiogram; bpm, beats per minute

**Table S2.** Genome sequencing of individual mouse lemurs.

| **Name** | **Sex** | **Age** | **SSS phenotype** | **Sequence coverage** | **Aligned reads** |
| --- | --- | --- | --- | --- | --- |
| 261BBA | M | 4.0 | Affected | 27X | 98.9% |
| 300AA | M | 3.4 | Affected | 40X | 99.5% |
| 300AB | M | 3.4 | Affected | 32X | 99.1% |
| 300AC | F | 2.5 | Affected | 37X | 99.5% |
| 241EA | F | 1.6 | Affected | 37X | 99.4% |
| 204FD | F | 1.5 | Affected | 112X | 99.5% |
| 261BB | F | 4.6 | Presumed carrier | 31X | 99.5% |
| 300A | F | 3.5 | Presumed carrier | 31X | 99.4% |
| 300ACA | M | 2.5 | Presumed carrier | 32X | 99.4% |
| 241E | F | 2.5 | Presumed carrier | 84X | 99.5% |
| 261BBC | F | 3.5 | Unaffected family member | 55X | 99.6% |
| 277AB | F | 4.9 | Outside family control | 30X | 99.4% |
| 208CBE | F | 3.7 | Outside family control | 48X | 99.4% |
| 288H | M | 1.6 | Outside family control | 56X | 99.3% |
| 223HCB | F | 2.4 | Outside family control | 54X | 99.0% |
| 245BAB | F | 3.3 | Outside family control | 32X | 99.5% |
| 374C | M | 1.5 | Outside family control | 31X | 99.2% |
| 283E | F | 2.3 | Outside family control | 34X | 99.1% |
| 352A | M | 3.5 | Outside family control | 29X | 99.0% |
| 245BAA | M | 3.4 | Outside family control | 42X | 98.9% |
| 275BBC | F | 1.4 | Outside family control | 34X | 99.3% |
| 277AH | F | 1.3 | Outside family control | 46X | 99.0% |
| 283BB | M | 2.4 | Outside family control | 88X | 98.9% |
| 316A | M | 3.9 | Outside family control | 108X | 99.2% |
| 277C | M | 3.9 | Outside family control | 201X | 99.4% |
| 283C | F | 4.9 | Outside family control | 293X | 99.2% |
| 288F | M | 4.5 | Outside family control | 129X | 99.4% |
| 244DBD | F | 2.4 | Outside family control | 149X | 99.5% |
| Antoine | M | 6.2 | Outside family control | 88X | 99.7% |
| Bernard | M | 5.4 | Outside family control | 67X | 99.6% |
| Inina | F | 2.3 | Outside family control | 67X | 99.7% |
| Martine | F | 11.8 | Outside family control | 75X | 99.7% |
| Nadine | F | 12.9 | Outside family control | 290X | 99.1% |
| Pascal | M | 3.5 | Outside family control | 67X | 99.6% |
| Stumpy | F | 5.4 | Outside family control | 86X | 99.7% |

**Table S3. Metadata for the 365 SSS-linked DNA sequence variants**

(Available upon request -- please contact authors for this file)

**Table S4.** Metadata for additional genes in the SSS mapped interval on Chromosome 7.

| Position | P1 | P2 | P3 | P4 | P5 | P6 |
| --- | --- | --- | --- | --- | --- | --- |
| Gene | POLR3B | TCP11L2 | NUAK1 | LOC105875605 | LOC105875604 | LOC105875644 |
| Gene product | RNA polymerase III subunit B | Promotes differentiation of muscle-derived satellite cells | NUAK family kinase 1 | Uncharacterized ncRNA, no human ortholog | Uncharacterized ncRNA, no human ortholog | Uncharacterized ncRNA, no human ortholog |
| Transcription | < | < | > | > | < | > |
| Lemur heart expression | - | - | Endothelial cell, Fibroblast, Myeloid cell | - | - | - |
| Mouse KO | Intestinal epithelium-specific KO results in decreased growth/survival during 1st postnatal week, decreased proliferation, abnormal epithelial architecture | NA | Global +/- show defective spatial memory consolidation, abnormal sensorimotor gating, abnormal social behavior | NA | NA | NA |
| Mouse IMPC | Embryonic and Prewaning lethality | Decreased BMD/lean body mass, increased body fat, abnormal locomotion | NA | NA | NA | NA |

| Position | P7 | P8 | P9 | P10 | P11 | P12 |
| --- | --- | --- | --- | --- | --- | --- |
| Gene | LOC105875643 | WASHC4 | C7H12orf45 | TXNRD1 | NFYB | HCFC2 |
| Gene product | Uncharacterized ncRNA, no human ortholog | Polymerizes actin at endosomes | Protein chaperone that binds to RNP complex | Thioredoxin reductase 1 | Nuclear transcription factor Y subunit beta | Host cell factor, initiates HSV infection |
| Transcription | < | < | < | < | > | < |
| Lemur heart expression | - | Endothelial cell, Fibroblast, Myeloid cell | Pericyte, Neutrophil | - | Endothelial cell, Fibroblast, Vascular smooth muscle cell | Endothelial cell, Fibroblast, Vascular smooth muscle cell |
| Mouse KO | NA | Mouse with human WASHC4(c.3056C>G) disrupts endo-lysosomal pathways, causes neurodegeneration, motor deficits | NA | Global -/- are embryonic lethal; Cardiomyocyte-specific mouse -/- appear healthy | NA | Decreased macrophage response to HSV |
| Mouse IMPC | NA | HOM: embryonic and | NA | NA | NA | NA |

Abbreviations: see Fig. 5 legend; Gene product, gene product function; Lemur heart expression, lemur heart cell types in which the given gene is highly and differentially expressed; "-", gene is not detected or differentially expressed; Mouse KO, mouse knockout phenotypes in published literature; Mouse IMPC, mouse knockout phenotypes determined by the International Mouse Phenotyping Consortium; BMD, bone mineral density; HSV, herpes simplex virus; HOM, homozygous; HET, heterozygous

**Movie S1. Video of lemur ECG recording**

